# Specification of bone marrow sinusoids requires TIE2-mediated positive feedback involving COUPTFII and VEGFR3

**DOI:** 10.64898/2026.05.17.725724

**Authors:** Xiao Li, Xiwen Jia, Zhiliang Sun, Taotao Li, Beibei Xu, Xudong Cao, Kai Ding, Yulong He

## Abstract

The bone marrow (BM) vascular network plays crucial roles in driving bone development and supporting hematopoiesis, yet the mechanisms governing its specialized architecture, particularly sinusoidal morphogenesis, remain inadequately characterized. We show in this study that TIE2 (*Tek*) was highly expressed by BM sinusoidal endothelial cells (SEC) and the endothelial *Tek* excision led to BM sinusoidal capillarization. Particularly, the BM sinusoids displayed thinner vessel diameter with the aberrant mural cell coverage in the *Tek* mutants. Mechanistically, TIE2 insufficiency led to a dramatic decrease of VEGFR3 in BM-SECs while its expression in hepatic sinusoids was not obviously altered. The RNA-seq analysis showed that GO terms enriched for the downregulated genes were related to the biological processes including sinusoidal development while pathways related to arterial ECs and angiogenesis were upregulated in the bone marrow of *Tek* mutants. The alteration of sinusoidal VEGFR3 expression occurred within 48 h after the induced endothelial deletion of *Tek*. Consistently, the defective BM sinusoidal formation was validated with the induced *Tek* deletion in VEGFR3^+^ SECs. The insufficiency of TIE2 ligand ANGPT1 also led to reduced sinusoidal VEGFR3, accompanied by similar BM sinusoidal defects. Furthermore, disruption of sinusoidal morphogenesis was observed in mutant mice with the endothelial excision of *Nr2f2* (COUP-TFII), displaying a decreased expression of BM sinusoidal TIE2 and VEGFR3. These findings suggest that ANGPT1/TIE2 and COUP-TFII form a reciprocal regulatory loop to coordinate BM sinusoidal specification via regulating VEGFR3.

## Introduction

Sinusoids are a specialized type of blood vessels distributed in several organs including bone marrow and liver (BM) [1, 2]. It has been reported that bone marrow sinusoidal vessels are derived from caveolin-1 (CAV1) positive type E capillaries, based on lineage-tracing studies using the *Apln-CreER* line [3, 4]. This involves the angiogenic growth followed by the remodeling of highly proliferating type E vessels to form the type H vessels (PECAM1^hi^) and the type L sinusoidal vessels (PECAM1^lo^; VEGFR3^hi^). The type H vessels are located in the metaphysis and the endosteal regions to regulate osteogenesis while the BM sinusoids are highly fenestrated for trafficking of hematopoietic cells into the circulation [3, 5].

Sinusoidal morphogenesis is tightly regulated involving a complex interplay of angiogenic factors and related signaling pathways, predominantly derived from endothelial cells as well as perivascular mesenchymal cells. In contrast to the early specialized hepatic sinusoids, bone marrow (BM) sinusoids form later as the primary site of adult hematopoiesis [6]. VEGFs, including VEGFA and VEGFC produced by mesenchymal stem cells (MSCs) and hematopoietic cells, are the primary drivers of angiogenesis and sinusoidal formation in bone marrow. They interact with VEGFR2 and VEGFR3 on sinusoidal endothelial cells (BM-SECs) to promote proliferation and maintain the fenestrated phenotype [7]. The VEGFR2-Notch signaling pathway is strongly associated with arteriolar differentiation, and its regulation is critical for the balance between the angiogenic vessels and sinusoids [3, 8]. Osteoblasts are important producers of vascular-remodeling factors including VEGF and ANGPT1, which directly influence the morphology and permeability of the adjacent sinusoids [9]. Angiopoietin receptor TIE2 is primarily expressed on vascular endothelial cells, mediating critical signals in the remodeling and maturation of vascular system [1, 10]. Particularly, TIE2 is crucial for the venous specification via the downstream PI3K/AKT-mediated pathway to regulate the protein stability of transcription factor COUP-TFII (Chicken ovalbumin upstream promoter-transcription factor II, gene name as *Nr2f2*) [11]. COUP-TFII is essential for the specification of venous EC identity by suppressing the Notch signal pathway [12, 13]. On the other hand, it has been reported that COUP-TFII drives the expression of TIE2 ligand angiopoietin-1 (ANGPT1) in pericytes [14]. Furthermore, COUP-TFII has been shown to regulate the expression of NRP2 and VEGFR3 [15]. VEGFR3 is mainly expressed by lymphatic endothelial cells (LECs) as well as a proportion of blood vascular endothelial cells (BECs) including sinusoidal vessels [7, 16, 17]. It has been shown recently that VEGFR3 is also required for the sinusoidal growth in both liver and bone marrow [7].

Single cell RNA-seq analysis revealed that a higher level of *Tek* transcripts was detected in bone marrow SECs compared with the type H capillary ECs in metaphysis [18]. Following the myelosuppressive stress by chemotherapy, TIE2 expression is rapidly upregulated on BM endothelial cells for the vascular regeneration [19, 20]. Despite its known role in vascular formation, the precise function of TIE2 in bone marrow sinusoidal specification remains to be fully elucidated. We show in this study that the induced deletion of endothelial *Tek* or *Angpt1* led to the defective formation of sinusoidal vessels in bone marrow. TIE2 or its ligand ANGPT1 insufficiency resulted in a dramatic decrease of VEGFR3 in SECs. Consistently, disruption of sinusoidal morphogenesis was also observed in mutant mice with the endothelial *Nr2f2* deletion, displaying a decreased expression of the sinusoidal TIE2 and VEGFR3. Findings from this study imply that the ANGPT1-TIE2 and COUPTFII form a positive feedback loop in the coordination of BM sinusoidal morphogenesis via regulating the expression of sinusoidal VEGFR3.

## Materials and methods

### Animal models

All animal experiments were performed in accordance with the institutional guidelines of Soochow University Animal Center. All the mice used in this study were housed in a SPF (specific pathogen free) animal facility with a 12/12 hours dark / light cycle and were free to food and water access. The *Tek^Flox/Flox^*and *Angpt2^Flox/Flox^* mouse lines were generated by the Model Animal Research Center of Nanjing University, as previously described [21]. The mouse line with *Angpt1* knockout first allele (*Angpt1^tm1a(KOMP)Wtsi^*) was established from EUCOMM embryonic stem cells (EPD0585-1A04), in which targeting cassette is recombined downstream of exon 2 (with exon 3 floxed). To obtain the *Angpt1^Flox/Flox^*mouse line, the lacZ reporter and neo-cassette were removed using the FLPeR mice as previously described [22]. The mouse line with *Angpt4* knockout first allele (*Angpt4^tm1a(KOMP)Wtsi^*, mice homozygous for the allele labelled as *Angpt4^-/-^*) was generated from EUCOMM embryonic stem cells (EPD0761-2F08), in which targeting cassette is recombined downstream of exon 3 (with exon 4 floxed). For the generation of ubiquitous or pan-endothelial cell (EC) specific gene deletion, the *Ubc-Cre^ERT2^*[23] and *Cdh5-Cre^ERT2^* [24] mouse lines were used. The *Vegfr3-Cre^ERT2^*mouse line, generated by the Model Animal Research Center of Nanjing University, was employed for the sinusoidal EC specific gene deletion. Briefly, the targeting vector, containing a His_6_ tag followed by a TAA stop codon and ires-creERT2 together with a FRT site flanked neo cassette, was introduced before the 3’ UTR of the *Flt4* gene. The targeted ES cell screening, mice generation and the removal of neo-cassette using the FLPeR mice were performed as previously described [22, 25]. In all the phenotype analysis, wild-type or heterozygous littermates were used as controls. For the genotyping of *Angpt1* knockout allele, the primers used were as follows: the forward primer (5’-TTCCAGTCAGTGCAGGTAGAGA -3’) and reverse primer (5’-CCATGATCCGAAGAGTGGAGT -3’), to amplify a 1202 bp fragment for the wild-type allele and a 475 bp fragment for the knockout allele. For the genotyping of *Angpt4* knockout allele, the primers used were as follows: the forward primer (5’-TTCCAGTCAGTGCAGGTAGAGA -3’) and reverse primer (5’-CCATGATCCGAAGAGTGGAGT -3’), to amplify a 336 bp fragment for the wild-type allele and a 499 bp fragment for the knockout allele. For the genotyping of *Tek* and *Angpt2* mouse lines, the primers used were as previously described [21]. *Nr2f2^Flox/Flox^* mice were kindly provided by Dr Tsai’s laboratory [26]. The genetic background of all mouse lines is on C57BL/6J or mixed C57BL/6J/SV129.

### Induced gene deletion

Induction of gene deletion was performed as previously described by tamoxifen treatment [11, 27]. Briefly, new-born pups were treated by three daily intragastric injections of tamoxifen (Sigma-Aldrich, T5648-5G) from the postnatal day 1 (P1, 60 μg/day) or P5 (80 μg/day). For the inducible gene deletion at adolescent or adult stages, the intraperitoneal injections of tamoxifen for five or seven consecutive days were performed from P21 (600□µg/day) or 8-week-old (1□mg/day). For the gene deletion at embryonic stages, pregnant mice were treated by the intraperitoneal injection of tamoxifen at E12.5–14.5 or E14.5–16.5 (1□mg / per mouse, for three consecutive days). The genotypes of the *Tek* mutants are as follows: *Tek^Flox/-^*;*Cdh5-Cre^ERT2^* labelled as *Tek^iKO;CDH5+^*, *Tek^Flox/-^*; *Vegfr3-Cre^ERT2^* labelled as *Tek^iKO;VEGFR3+^* and the corresponding littermate control mice (labelled as *control*) are *Tek^Flox/+^*; *Cdh5- Cre^ERT2^*, *Tek^Flox/+;^* ; *Vegfr3-*Cre*^ERT2^* or *Tek^Flox/+.^*. The genotypes of the *Angpts* knockouts are labelled as follows: *Angpt1^iKO;UBC+^*(*Angpt1^Flox/-^*; *Ubc-Cre^ERT2^*), *Angpt2^iKO;CDH5+^*(*Angpt2^Flox/-^*; *Cdh5-Cre^ERT2^*). The corresponding littermate control mice are labeled as control, including *Angpt1^Flox/+^*(or *Angpt1^Flox/+^*; *Ubc-Cre^ERT2^*), *Angpt2^Flox/+^* (or *Angpt2^Flox/+^*; *Cdh5-Cre^ERT2^*), and *Angpt4^+/+^* mice. The genotypes of the *Nr2f2* knockout and control mice are labelled as follows: *Nr2f2^iKO;CDH5+^*(*Nr2f2^FloxFlox^*; *Cdh5-Cre^ERT2^*), and *Nr2f2^Flox/Flox^*used as the control. Tissues were collected for analysis at the specified stages. Histological analysis of femur bone and retina tissues was performed according to the previous reports [5, 28]. The retinal vascularization index was quantified as the ratio of vascularized area to total retinal area as previously published [11].

### Immunostaining

For immunostaining, the whole retina tissues, femur bone sections (70 µm) or liver sections (15 µm) were blocked with 5% (v/v) donkey serum in PBS-TX (0.3% Triton X-100), and then incubated the primary antibodies at 4□°C overnight. The following antibodies were used: Goat-anti-mouse TIE2 (R&D AF762), Rat-anti-mouse endomucin (EMCN, eBioscience 14-5851), rat-anti-mouse PECAM1 (BD 553370), Hamster-anti-mouse PECAM-1 (Thermo Scientific MAB13982), Rabbit-anti-mouse NG2 (Millipore AB5320), Goat-anti-mouse VE-Cadherin (CDH5, R&D AF1002), Goat-anti-mouse VEGFR3 (R&D AF743), Rabbit-anti-mouse Caveolin-1 (CAV1, Cell Signaling 3238), Goat-anti-mouse VEGFR2 (R&D AF644), Rabbit-anti-mouse LYVE1 (Abcam ab14917), eFluor 660-αSMA (eBioscience 50976082). Appropriate Alexa488, Alexa546 (Invitrogen), Cy3- or Cy5- (Jackson) conjugated secondary antibodies were applied for detection. Fluorescently labeled samples were mounted and analyzed with a confocal microscope (Olympus Flueview 1000 or 3000). For comparison, the parameters were kept consistent for all the confocal microscopic imaging in this study.

### Quantification of sinusoidal vessel

For the quantification of sinusoidal vessel diameter in the bone marrow, images were taken from the mid-diaphyseal regions (with×20 magnification). Five vertical lines were evenly laid on the images, and the diameters of sinusoidal vessels crossed with these lines were measured and analyzed using Image Pro Plus (Media Cybernetics, Inc., Bethesda, MD). Furthermore, quantifications of fluorescence intensity in BM vessels positive the EC markers (including EMCN, PECAM1 or VEGFR3) and the VEGFR3^−^/CAV1^+^ areas were performed using Image Pro Plus.

### Western blot

Bone marrows flushed from the mouse femur bones, liver or lung tissues were lysed in NP-40 lysis buffer (Beyotime P0013F) supplemented with protease inhibitor cocktail (complete Mini, Roche 04693124001), phosphatase inhibitor cocktail (PhosSTOP, Roche 04906837001), 10 mmol/L NaF and 1 mmol/L PMSF. Protein concentration was determined using the BCA protein assay kit (PIERCE), and equal amounts of protein were used for analysis. Briefly, after the protein transfer from gels to PVDF membranes (IPVH00010, Millipore) and the antibody incubation, images were acquired by the chemiluminescent detection method (NEL105001EA, PerkinElmer) using X-ray film (XBT, 6535876, Carestream). For images acquired using X-ray film, the protein markers were manually marked on the films overlapped with the PVDF membranes with the prestained protein ladder (26616, ThermoFisher Scientific). The entire blot was further washed and re-probed with antibodies for beta-actin (ACTB) as loading controls. The following antibodies were used in this study, including Goat-anti-mouse TIE2 (R&D AF762), Goat-anti-mouse VEGFR3 (R&D AF743), Rabbit-anti-mouse CAV1 (Cell Signaling 3238), goat polyclonal anti-mouse ANGPT4 (R&D AF738), mouse monoclonal to beta-actin antibody (Santa Cruz sc-47778), goat anti-mouse IgG, HRP conjugated (Fcmacs Biotech FMS-MS01), bovine anti-goat IgG, HRP conjugated (Jackson ImmunoResearch Labs 805-035-180), goat anti-rabbit IgG, HRP conjugated (R&D Systems HAF008).

### RNA sequencing analysis

For the bulk RNA-seq analysis of femur tissues, freshly isolated femurs were cleaned to remove adherent muscles, periosteum, and cartilage tissue, and then snap-frozen in liquid nitrogen. Total RNA was isolated from the femur bone tissues using Trizol (Invitrogen) according to the manufacturer’s instructions. RNA-seq procedures were performed as previously reported [29]. Briefly, the sequencing was carried out on DNBSEQ platform (BGI-Shenzhen, China) with pairedlzlend 150 bp reads, and data were processed using the predefined analysis pipeline. Differential expression analysis was performed with using the R package DESeq2 (v 1.42.1). Functional enrichment analysis was conducted using clusterProfiler (v4.10.1) together with org.Mm.eg.db (v 3.18.0). The hallmark gene sets used in the GSEA (Gene Set Enrichment Analysis), including the hallmarks for sinusoidal, arterial and other type endothelial cells, are mainly defined according to the single cell transcriptome analysis of murine bone marrow, the molecular signatures database (mh.all.v2024.1.Mm.symbols) and the related GO terms [29–32]. The raw and processed data of RNA-seq analysis will be deposited in GEO upon publication.

### Quantitative real-time RT-PCR

cDNA was synthesized using the RevertAid First Strand cDNA Synthesis Kit (Thermo Scientific K1622). Real-time quantitative PCR was performed using a SYBR Premix Ex Taq kit (Takara RR420A) on the ABI QuantStudio6 system. The primers used were as follows: *Gapdh*: 5’-GGTGAAGGTCGGTGTGAACG-3’, 5’-CTCGCTCCTGGAAGATGGTG-3’; *Tek*: 5′- GATTTTGGATTGTCCCGAGGTCAAG-3′, 5′- CACCAATATATCTGGGCAAATGATGG-3′; *Angpt1*: 5′-CACATAGGGTGCAGCAACCA -3′, 5 ′ - CGTCGTGTTCTGGAAGAATGA ′; *Angpt2*: 5’- ATCCAACAGAATGTGGTGCAGAA-3’, 5’- AGCTCGAGTCTTGTCGTCTGG-3’. The transcripts of endothelial markers were normalized against *Gapdh*, and the relative expression level of every gene in mutant mice was normalized against that of littermate controls.

### Statistical analysis

For the 2-group comparison, the unpaired t test was performed with Welch correction if data passed the D’Agostino-Pearson or Shapiro-Wilk normality test, or the unpaired nonparametric Mann-Whitney U test was applied using GraphPad Prism v8.0. Data are expressed as mean ± SD. All statistical tests were 2-sided.

## Results

### Disruption of BM sinusoidal formation after the induced endothelial deletion of Tek

To investigate the role of TIE2 in bone marrow sinusoidal formation, we generated an inducible endothelial cell-specific *Tek* knockout mouse model (*Tek^Flox/-^*;*Cdh5-Cre^ERT2^*, named *Tek^iKO;CDH5+^*), and *Tek^Flox/+^*; *Cdh5-Cre^ERT2^*littermates served as controls. The scheme for *Tek* deletion by the intragastric administration of tamoxifen is shown in **Fig. 1A**. The efficiency of induced *Tek* deletion was examined by the immunostaining for EMCN and TIE2 (**Fig. 1B**), western blot analysis (**Fig. 1C-D**) and quantitative RT-PCR (**Fig. 1E**). TIE2 insufficiency was also confirmed by the suppression of retinal vascularization in the *Tek^iKO;CDH5+^*mice as shown in **Supplemental Fig. 1A-B**. The *Tek* mutant mice showed lethality starting approximately from the postnatal day 12 (P12). Detailed analysis of the survival rate revealed that all the mutant mice died before the age of 3-week-old (**Supplemental Fig. 1C**). There was also a significant decrease in the body weight and femur length of the *Tek^iKO;CDH5+^*mice at P11 (**Fig. 1F-G**; **Supplemental Fig. 1D-E**).

**Fig. 1.**
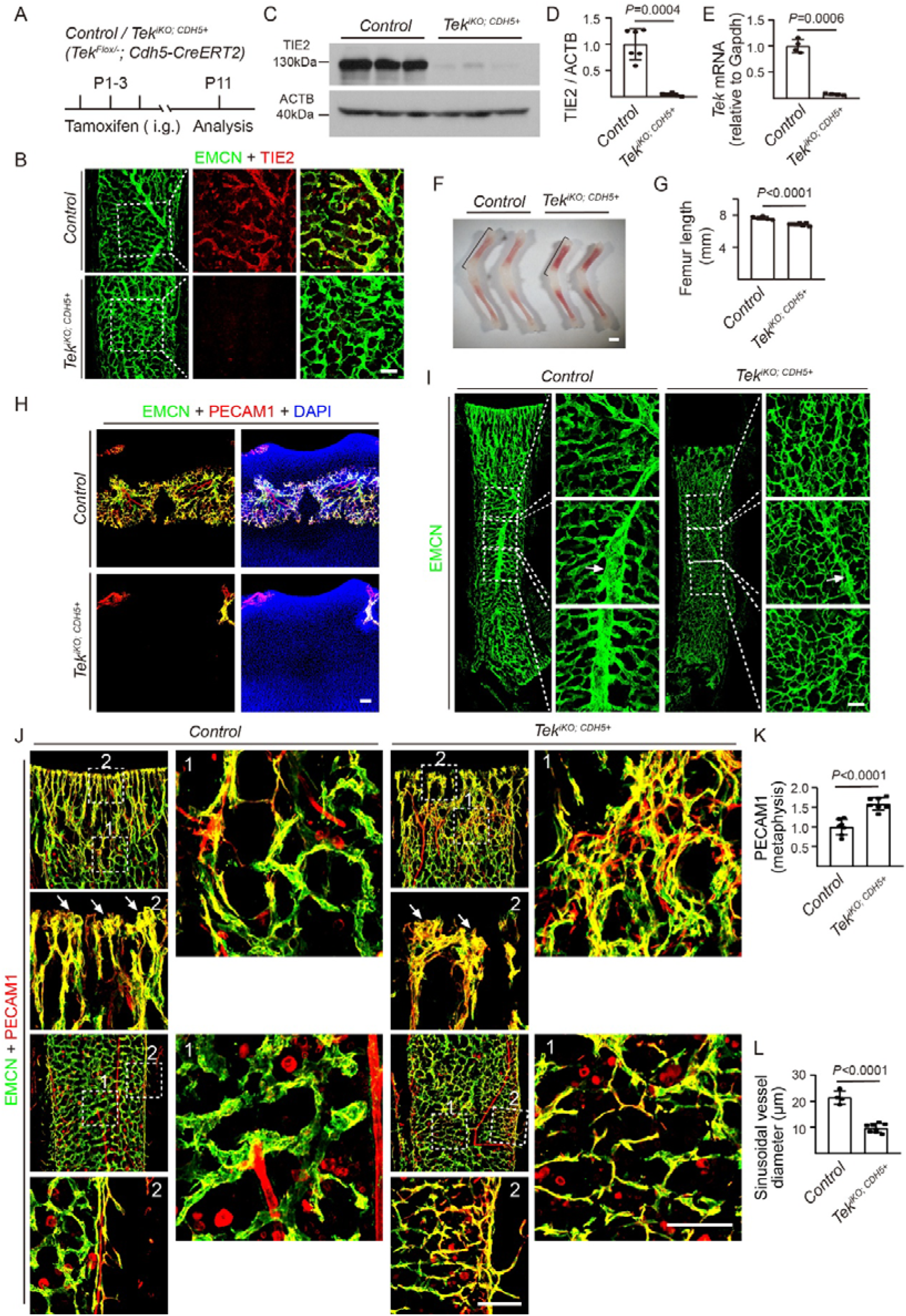
Endothelial TIE2 insufficiency disrupts bone marrow sinusoidal vessel formation. **A**. Tamoxifen intragastric (i.g.) administration and analysis scheme. **B**. Immunostaining for EMCN and TIE2 to analyze *Tek* deletion in bone marrow endothelial cells at postnatal day 11 (P11). **C** and **D**. Western blot analysis of TIE2 in bone marrow from *Tek^iKO;CDH5+^*and littermate control mice at P11. Quantification of TIE2 protein levels were normalized to beta-actin (TIE2/beta-actin, *Control*: 1.00 ± 0.29, n=6; *Tek^iKO;CDH5+^*: 0.04 ± 0.03, n=6, *P*=0.0004). **E**. Analysis of *Tek* deletion efficiency by qPCR (*Control*: 1.00 ± 0.13, n=4; *Tek^iKO;CDH5+^*: 0.08 ± 0.01, n=4; *P*=0.0006). **F-G**. Representative images and quantification of femur length from P11 *Tek^iKO;CDH5+^* and littermate control mice (*Control*: 7.64 ± 0.19 mm, n=6; *Tek^iKO;CDH5+^*: 6.84 ± 0.15 mm, n=7, *P*<0.0001). **H-J**. Analysis of bone marrow vascular network from *Tek^iKO;CDH5+^* and littermate control mice (P11) by immunostaining for EMCN and PECAM1, including secondary ossification center (SOC) in the epiphysis (H), BM central veins (arrows, I), vascular arch structures in the metaphysis, and the diaphysis (arrows, J). **K**. Quantification of angiogenesis at the metaphyseal-diaphyseal regions (transition zones) based on PECAM1 fluorescence intensity (*Control*: 1.00 ± 0.20, n=7; *Tek^iKO;CDH5+^*: 1.59 ± 0.16, n=7, *P*<0.0001). **L**. Quantitative analysis of sinusoidal vessel diameter in the diaphysis (*Control*: 21.40 ± 2.54 µm, n=4; *Tek^iKO;CDH5+^*: 9.73 ± 1.64 µm, n=8, *P*<0.0001). Scale bar: 500 μm in **F**, and 100 μm in **B** and **H-J**.

By the histological analysis of femur tissues from the *Tek* mutant and control mice at P11, we found that TIE2 insufficiency delayed vascularization of the secondary ossification center in the epiphysis **(Fig. 1H**, **Supplemental Fig. 2A-C**). Consistent with the previous findings about TIE2 in venous specification [11], we observed that the formation of bone marrow central veins was suppressed in the *Tek^iKO;CDH5+^* mice (P11, **Fig. 1I**). As shown in **Fig. 1J-L**, the process of BM vascular morphogenesis was disrupted in the femur tissues of *Tek* mutants, showing a significant decrease of sinusoidal diameter in the diaphysis (**Fig. 1J and L**). The BM vascularization, including the vascular arch structures at the distal edge of the metaphysis and the sinusoidal maturation in the diaphysis, was disrupted due to the abnormal increase of angiogenesis (**Fig. 1J and K**). The abnormal sinusoidal capillarization in the diaphysis was accompanied by the aberrant coverage of perivascular mural cells (NG2⁺) in the *Tek^iKO;CDH5+^* mice. However, there was no obvious alteration of endothelial adherens junctions as shown by the immunostaining for CDH5 after the endothelial deletion of *Tek* (**Supplemental Fig. 1F-G**).

### VEGFR3 as a downstream effector of TIE2 in sinusoidal morphogenesis

VEGFR3 mediates an essential signaling pathway for lymphatic growth and it is also expressed by a proportion of blood vascular endothelial cells including sinusoidal ECs [16, 17, 33]. In contrast, caveolin-1 (CAV1) is primarily expressed in arteries and blood capillaries, but low in BM sinusoidal vessels [3, 34]. We found in this study that TIE2 was mainly expressed in EMCN⁺ sinusoids and relatively low in CAV1⁺ vessels (**Fig. 2B**). Interestingly, the induced deletion of endothelial *Tek* resulted in a dramatic decrease of VEGFR3 in BM sinusoidal vessels while CAV1 was significantly upregulated (**Fig. 2A**, **C**). The alteration of VEGFR3 and CAV1 was further confirmed by western blot analysis of the femur tissues of *Tek^iKO;CDH5+^* mice compared with littermate controls (**Fig. 2D**, **E**).

**Fig. 2.**
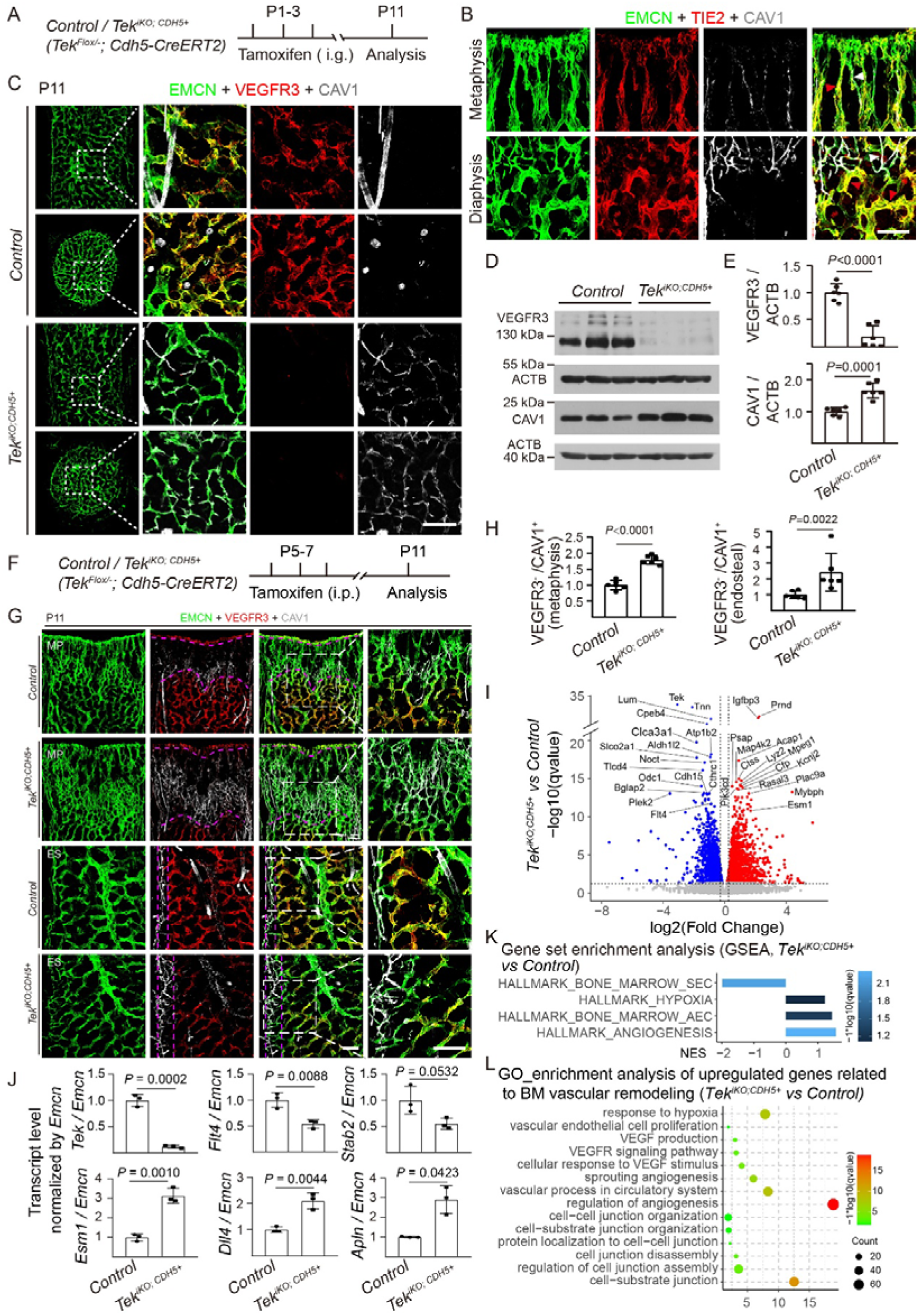
TIE2 insufficiency leads to the downregulation of sinusoidal VEGFR3. **A**. Tamoxifen intragastric (i.g.) administration and analysis scheme. **B**. Expression analysis of TIE2 and CAV1 in BM blood vessels of metaphysis and diaphysis from wild-type mice (P15). White arrowheads indicate CAV1^+^ vessels, red arrowheads indicate CAV1^−^ vessels. **C**. Immunostaining for VEGFR3 and CAV1 in bone marrow sinusoidal vessels from *Tek^iKO;CDH5+^*and littermate control mice at P11. **D** and **E**. Western blot analysis of VEGFR3 and CAV1 in bone marrow from *Tek^iKO;CDH5+^* and littermate controls (VEGFR3/beta-actin, *Control*: 1.00 ± 0.16, n=6; *Tek^iKO;CDH5+^*: 0.18 ± 0.20, n=6, *P*<0.0001; CAV1/beta-actin, *Control*: 1.00 ± 0.13, n=6; *Tek^iKO;CDH5+^* mice: 1.66 ± 0.24, n=6, *P*=0.0001). **F**-**G**. Tamoxifen treatment and analysis scheme, and immunostaining analysis of sinusoidal vessels for EMCN, VEGFR3 and CAV1 in metaphysis (MP) and diaphysis (endosteum, ES) from *Tek^iKO;CDH5+^* and littermate control mice at P11. **H.** Quantification of VEGFR3^−^ / CAV1^+^ areas in metaphysis (*Control*: 1.00 ± 0.15, n=6; *Tek^iKO;CDH5+^*: 1.79 ± 0.14, n=6; *P*<0.0001) and endosteum (*Control*: 1.00 ± 0.24, n=6; *Tek^iKO;CDH5+^*: 2.42 ± 1.19, n=6, *P*=0.0022). **I**-**L**. RNA-seq analysis of the femur tissues from *Tek^iKO;CDH5+^* and control mice at P11. Volcano plot visualization of the differentially expressed genes (DEGs, P-value ≤ 0.05, |Log2Foldchange| > 0) (I). Quantification of sinusoidal (*Tek*, *Flt4*, *Stab2*) and angiogenic gene expression (*Esm1*, *Dll4*, *Apln*), normalized by *Emcn* (J). GSEA analysis showed that the genes involved in sinusoidal development were significantly downregulated and the upregulated genes were related to hypoxia, BM arterial ECs and angiogenesis after the endothelial TIE2 insufficiency (K). Gene Ontology (GO) analysis further confirmed that genes related to hypoxia, angiogenesis and endothelial cell junction remodeling were significantly upregulated (L). Scale bar: 100 µm.

Vascularization of the murine femur was initiated at approximately E14.5 [3, 35]. To dissect the role of TIE2 in BM sinusoidal development, we performed the *Tek* gene deletion at embryonic day 14.5 (E14.5) and embryos were collected for analysis at E18.5. Consistently, we found that the BM sinusoidal VEGFR3 decreased while there was an increase in CAV1 expression (**Supplemental Fig. 3A, B**). In addition, to minimize the effect of TIE2 on the BM central vein formation, we also performed the *Tek* deletion at a later stage starting from the postnatal day 5 (P5). As shown in **Fig. 2F-H**, the BM central veins were detected, but the sinusoidal vessels became thinner with the decreased expression of VEGFR3 in the *Tek^iKO;CDH5+^* mice. Consistently, the transition zone between the metaphysis and diaphysis was expanded, and there was also an increase of VEGFR3^-^ and CAV1^+^ area in the endosteum of *Tek* mutants compared with littermate controls.

### Alteration of sinusoidal transcriptome upon the endothelial TIE2 insufficiency

To explore the mechanism underlying TIE2 in BM sinusoidal morphogenesis, we performed the RNA sequencing (RNA-seq) analysis of femur tissues to examine the alteration of transcriptome upon the induced deletion of endothelial *Tek*. TIE2 insufficiency led to a change in the overall transcriptional profile (**Fig. 2I**), including the suppression of gene expression related to the sinusoidal development, such as *Flt4* (encoding VEGFR3) and *Stab2*, while the arterial /angiogenic gene were upregulated including *Esm1*, *Dll4* and *Apln* (**Fig. 2J**). Consistently, pathways related to sinusoidal development were downregulated, while the hypoxia, angiogenesis and arterial EC related pathways were upregulated by the GSEA analysis using the custom gene sets as described above (**Fig. 2K-L**).

To explore the direct response upon endothelial TIE2 insufficiency, we performed the phenotype and molecular analysis of femur tissues from the *Tek* mutants and control mice 24 to 96 hours after the induced gene deletion (**Fig. 3A-D**; **Supplemental Fig. 3C-E**). While the decrease of TIE2 protein was detected 24 hours after the induced deletion of endothelial *Tek* (**Fig. 3B**), there was no obvious change with the sinusoidal VEGFR3 (**Fig. 3C**). The decrease of VEGFR3 protein was readily detectable 48 hours later, as shown by the immunostaining for VEGFR3 and EMCN (**Fig. 3D**). The alteration of sinusoidal morphology was also confirmed at 96 hours after the induced deletion of endothelial *Tek*, accompanied by the decrease of sinusoidal VEGFR3 and the increase of CAV1 (**Supplemental Fig. 3C-E**). The decrease of *Flt4* transcripts (VEGFR3) was further confirmed by the RNA-seq analysis of femur tissues 48 hours after the induced deletion of endothelial *Tek* (*Tek^iKO;CDH5+^*), as shown in **Fig. 3E-H**. The altered expression of sinusoidal EC genes was shown by the volcano plot and heatmap (**Fig. 3E-F**). Consistently, pathways related to BM sinusoidal development were downregulated in the *Tek^iKO;CDH5+^* mutants by the GSEA analysis (**Fig. 3G**). Shown in **Fig. 3H** was the quantification of the transcript levels of sinusoidal EC genes (e.g. *Flt4* and *Stab2*) and angiogenic /arterial EC genes (e.g. *Esm1*, *Dll4* and *Apln*) normalized by *Emcn* (48 h).

**Fig. 3.**
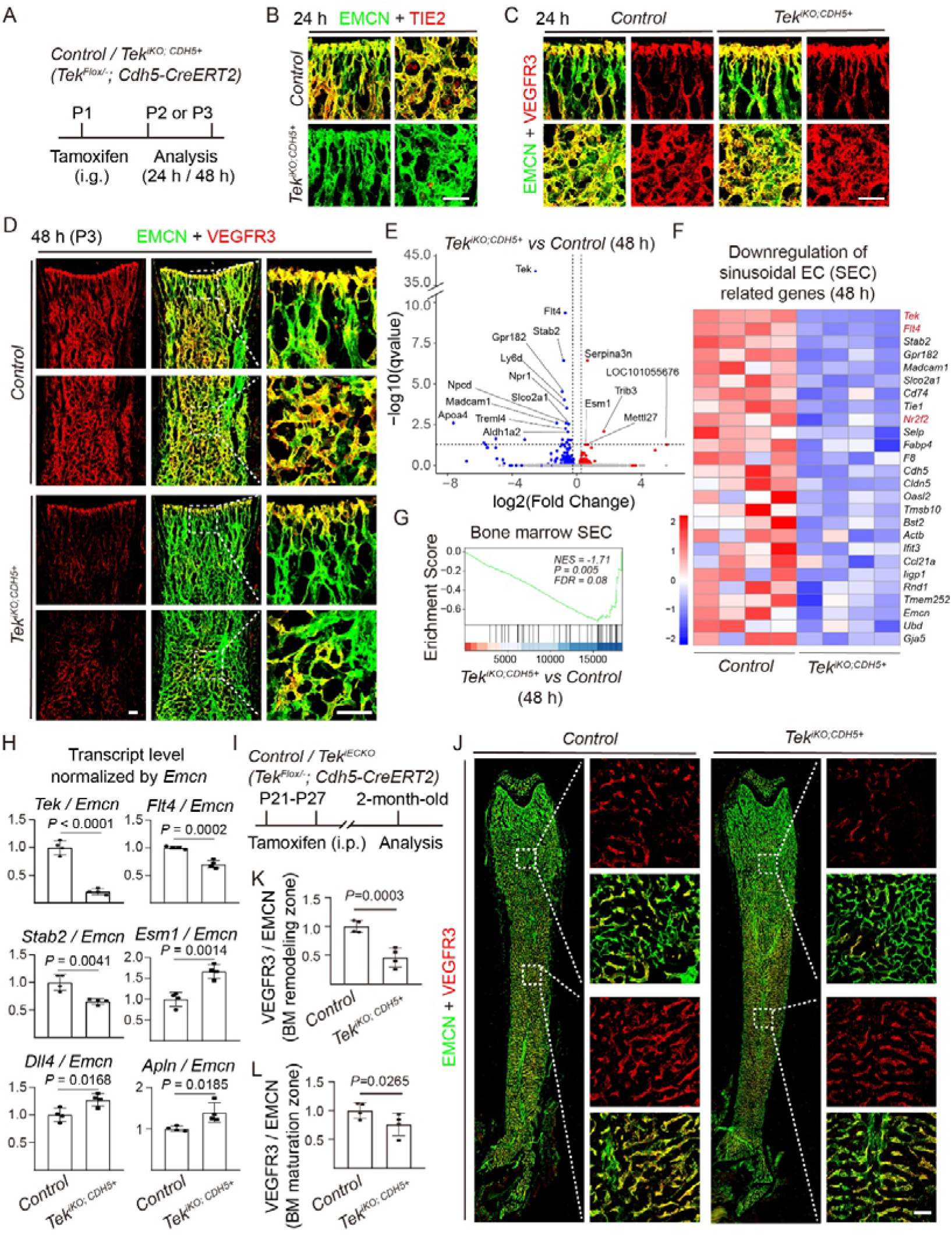
Requirement of TIE2 for the BM sinusoidal EC transcriptome and its maintenance. **A**. Tamoxifen intraperitoneal (i.g.) administration and analysis scheme. **B**. Analysis of TIE2 expression by immunostaining for EMCN and TIE2 in bone marrow vessels at 24 hours (h) after the induced *Tek* deletion. **C** and **D**. Immunostaining analysis of bone marrow sinusoidal vessels for VEGFR3 and EMCN in *Tek^iKO;CDH5+^* and littermate control mice at 24 h (C) and 48 h (D). **E**-**H**. RNA-seq analysis of the femur tissues from *Tek^iKO;CDH5+^* and control mice at 48 h (P3). Volcano plot showing differentially regulated genes (DEGs, P-value ≤ 0.05, |Log2Foldchange| > 0, E). Heatmap showing subsets of downregulated sinusoidal endothelial genes (*P*-value ≤ 0.05, F). GSEA analysis showed that the genes involved in sinusoidal development were significantly downregulated (G). Quantification of sinusoidal (*Tek*, *Flt4*, *Stab2*) and angiogenic gene expression (*Esm1*, *Dll4*, *Apln*), normalized by *Emcn*, in the femur of *Tek^iKO;CDH5+^* and littermate control mice at 48 h (H). **I**. Tamoxifen intraperitoneal (i.p.) administration and analysis scheme. **J**. Immunostaining for BM sinusoidal VEGFR3, CAV1 and EMCN after the induced excision of endothelial *Tek* at adult stages (2-month-old). **K** and **L**. Quantification of VEGFR3 fluorescence intensity relative to EMCN in the transition zones (at the metaphyseal-diaphyseal junction, *Control*: 1.00 ± 0.13, n=5; *Tek^iKO;CDH5+^*: 0.46 ± 0.15, n=5, *P*=0.0003; maturation zone of diaphysis, *Control*: 1.00 ± 0.12, n=5; *Tek^iKO;CDH5+^*: 0.75 ± 0.17, n=5, *P*=0.0265). Scale bar: 100 µm.

To further assess the role of TIE2 in bone marrow sinusoidal vessel maintenance, the induced gene excision was performed at 3 or 8 weeks of age, and tissues were collected for analysis four weeks later (**Fig. 3I, Supplemental Fig. 4A**). As shown in **Fig. 3J**, the decrease of sinusoidal VEGFR3 was detectable in the BM transition zone (the metaphyseal-diaphyseal interface). However, there were no obvious changes detected with the morphology of mature sinusoids in the diaphysis with a slight decrease of sinusoidal VEGFR3 (**Fig. 3J-L**). Consistently, the induced deletion of endothelial *Tek* at the adult stage produced a similar phenotype, with an alteration of sinusoidal VEGFR3 and vessel morphology in the transition zone of metaphysis when analyzed four weeks later (**Supplemental Fig. 4A-D**). In contrast, there was no obvious change with hepatic sinusoidal expression of VEGFR3 in *Tek^iKO;CDH5+^* mutants compared with the littermate controls at the embryonic (**Supplemental Fig. 5A-B**) or the postnatal stages (**Supplemental Fig. 5C-D**). Furthermore, the induced deletion of endothelial *Tek* did not alter the expression of VEGFR2 in BM sinusoidal ECs (**Supplemental Fig. 5E-F**).

### Induced deletion of *Tek* in VEGFR3^+^ sinusoidal endothelial cells

VEGFR3 is mainly expressed by the sinusoidal ECs in the BM vascular network, with a low level in venous ECs but not in arterial vessels [36].

To further investigate the role of TIE2 in BM sinusoidal ECs, we employed *Vegfr3-Cre^ERT2^* deleter line to generate mutant mice with *Tek* deletion in sinusoidal vessel (named as *Tek^iKO;VEGFR3+^*), and littermates were used as controls. The scheme for *Tek* deletion by the intragastric administration of tamoxifen is shown in **Fig. 4A**. The efficient deletion of *Tek* in sinusoidal ECs was examined by the immunostaining analysis for TIE2 and EMCN with both liver and femur tissues from the *Tek* mutant and control mice (**Fig. 4B, D**), and also by western blot analysis (**Fig. 4C**). In contrast to the pan-endothelial *Tek* mutant mice, the sinusoidal TIE2 insufficiency did not produce obvious effect on femur length at the postnatal day 11 (P11; **Fig. 4E-F**), and the BM central veins were still detectable in the *Tek^iKO;VEGFR3+^* mice (P11, **Fig. 4G**).

**Fig. 4.**
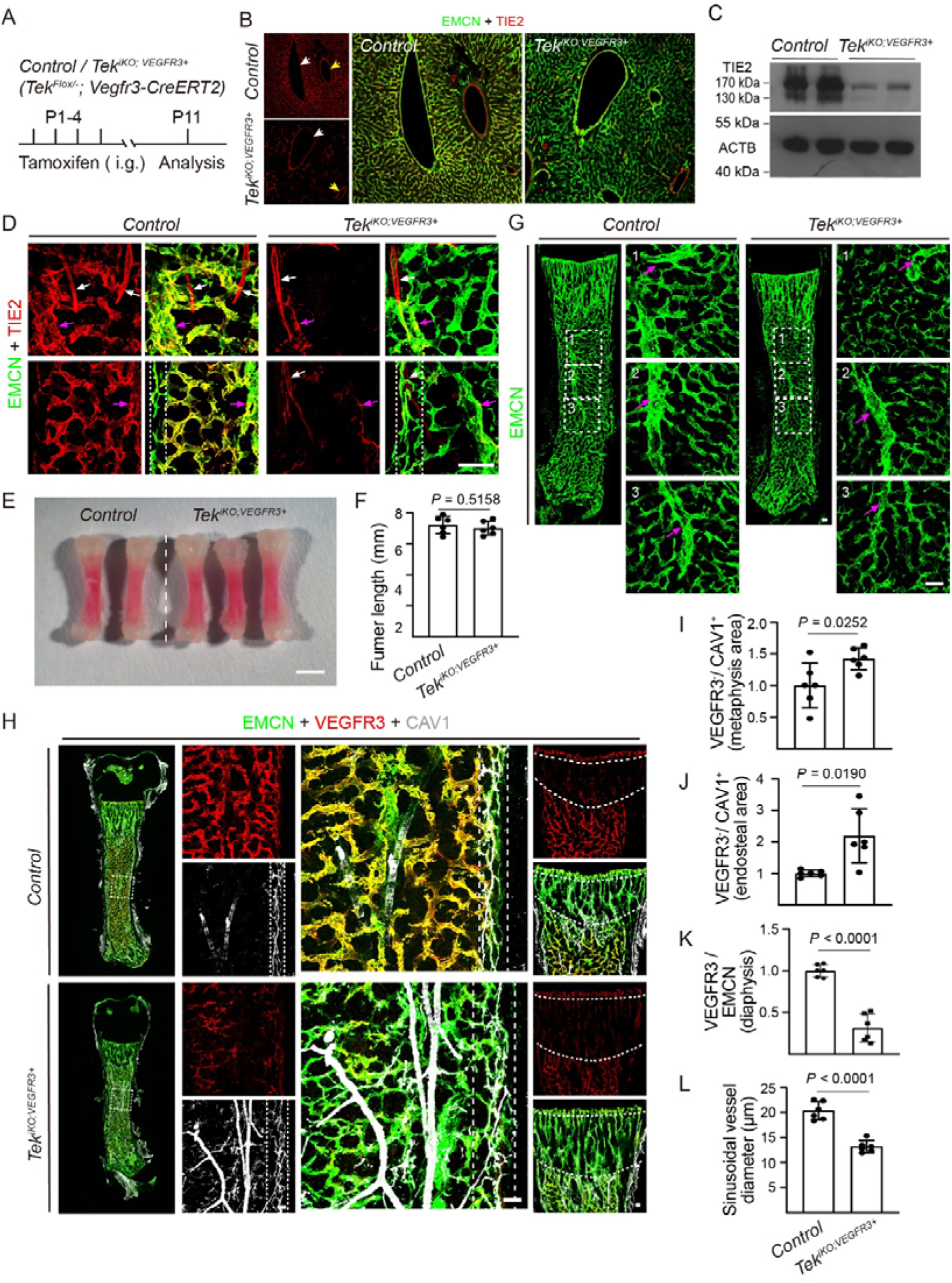
Defective BM sinusoidal formation upon the induced deletion of *Tek* in VEGFR3+ SECs. **A**. Tamoxifen intragastric (i.g.) administration and analysis scheme. **B**. Analysis of sinusoidal *Tek* deletion in the liver tissues of *Tek^iKO;VEGFR3+^* and littermate control mice at P11, employing a sinusoidal EC Cre deleter (*Vegfr3-CreERT2*). Yellow arrows indicate arteries, and white arrows indicate veins. **C**. Western blot analysis of TIE2 in liver tissues at P11. **D**. Analysis of sinusoidal *Tek* deletion by immunostaining for EMCN and TIE2 in bone marrow vessels at P11. White arrows indicate arteries, and pink arrows indicate central vein. **E**. Representative images and quantification of femur length from *Tek^iKO;VEGFR3+^*and littermate control mice (P11; *Control*: 7.22 ± 0.51 mm, n=6; *Tek^iKO;VEGFR3+^*: 7.02 ± 0.42 mm, n=6, *P*=0.5158). **G**. Analysis of BM vascular network by the immunostaining for EMCN in femur tissues from *Tek^iKO;VEGFR3+^*and littermate control mice at P11. Pink arrows indicate the normal central vein in the bone marrow. **H.** Immunostaining for BM sinusoidal VEGFR3, CAV1 and EMCN after the induced excision of sinusoidal *Tek* in femur tissues from *Tek^iKO;VEGFR3+^* and littermate control mice at P11. **I.** Quantification of the VEGFR3^−^ / CAV1^+^ areas in BM transition zone (*Control*: 1.00 ± 0.32, n=6; *Tek^iKO;VEGFR3+^*: 1.42 ± 0.16, n=6; *P*=0.0252). **J.** Quantification of the VEGFR3^−^ / CAV1^+^ areas in endosteum (*Control*: 1.00 ± 0.10, n=6; *Tek^iKO;VEGFR3+^*: 2.20 ± 0.79, n=6, *P*=0.0190). **K.** Quantification of VEGFR3 fluorescence intensity relative to EMCN in diaphysis (*Control*: 1.00 ± 0.07, n=6; *Tek^iKO;VEGFR3+^*: 0.31 ± 0.15, n=6, *P*<0.0001). **L**. Quantification of sinusoidal vascular diameter in the diaphysis (*Control*: 20.33 ± 1.68 µm, n=6; *Tek^iKO;VEGFR3+^*: 13.16 ± 1.09 µm, n=6, *P*<0.0001). Scale bar:100 μm.

Consistently, there was a marked increase with the VEGFR3^-^ / CAV1^+^ area at the BM transition zone and also endosteal region of the *Tek^iKO;VEGFR3+^* mice in comparison to that of controls (**Fig. 4H-J**). The induced deletion of sinusoidal *Tek* also resulted in a decrease of VEGFR3 in sinusoidal vessels while CAV1 was significantly upregulated (**Fig. 4H** and **K**). Similar morphological defects with the sinusoidal development were detected in the *Tek^iKO;VEGFR3+^* mutants, showing a significant decrease of sinusoidal vessel diameter in the diaphysis (**Fig. 4H** and **L**).

### Requirement of ANGPT1 but not ANGPT2/4 in BM sinusoidal formation

To further investigate the roles of the TIE2 ligands in bone marrow sinusoidal vessels, we employed genetically modified mouse models targeting *Angpt1* (*Angpt1^Flox/-^*;*Ubc-Cre^ERT2^*, named as *Angpt1^iKO;UBC+^*), *Angpt2* (*Angpt2^Flox/-^*; *Cdh5-Cre^ERT2^*, named as *Angpt2^iKO;CDH5+^*), and *Angpt4* (*Angpt4^-/-^*). The scheme for *Angpt1* deletion by the intragastric administration of tamoxifen is shown in **Fig. 5A**. The deletion efficiency of *Angpt1* was examined by the quantitative RT-PCR (**Fig. 5B**). Consistent with the above observations with *Tek* mutants, the induced deletion of *Angpt1* resulted in an obvious decrease of VEGFR3 in sinusoidal vessels while CAV1 was upregulated by the immunostaining (**Fig. 5C**). Shown in **Fig. 5D** is the VEGFR3 fluorescence intensity normalized by EMCN in the diaphysis. The alteration of VEGFR3 was further confirmed by the Western blot analysis of the femur tissues of *Angpt1^iKO;UBC+^* mice compared with littermate controls (**Fig. 5E**). The alteration of sinusoidal formation was detected, showing a significant decrease of sinusoidal vessel diameter in the diaphysis (**Fig. 5C** and **F**).

**Fig. 5.**
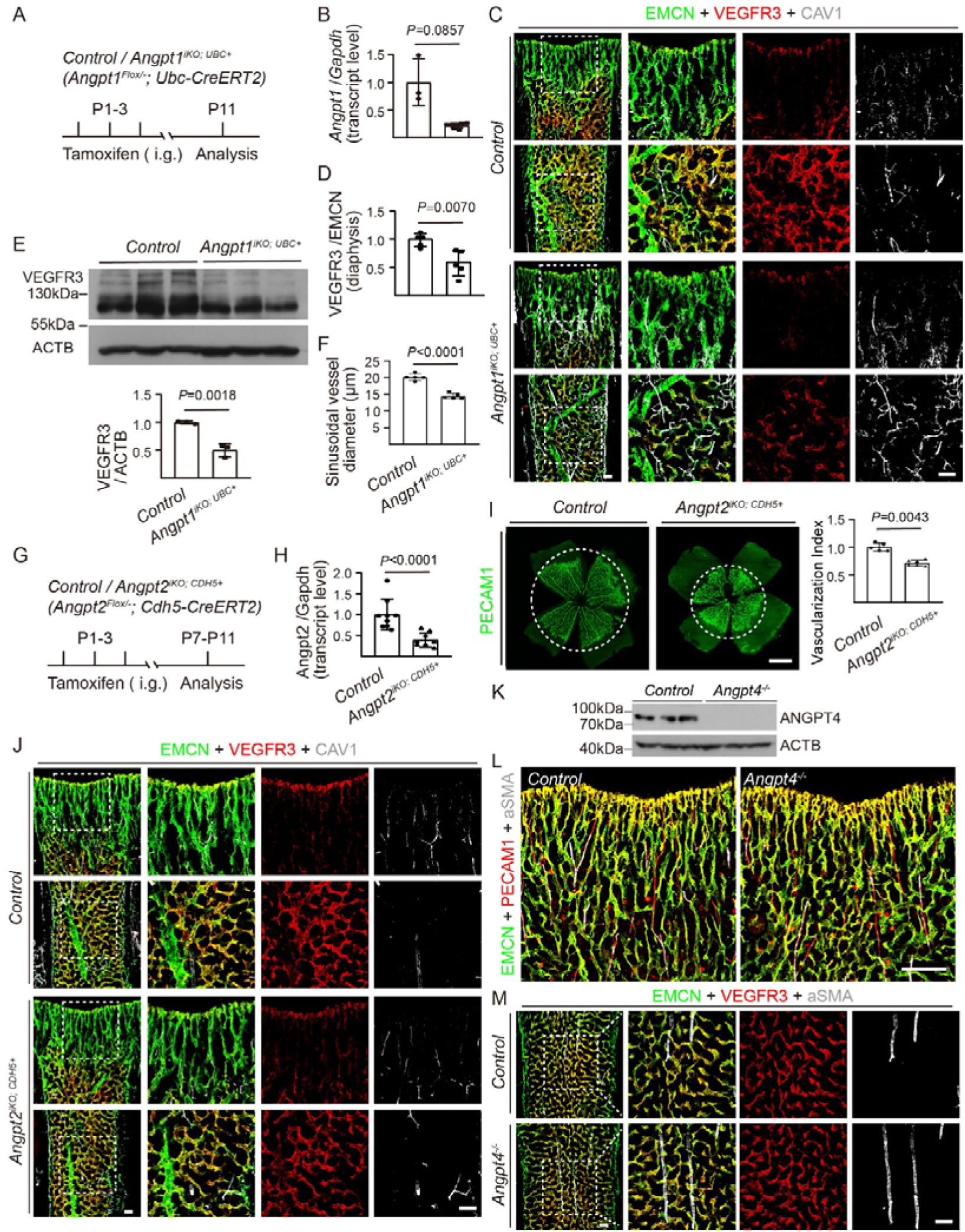
ANGPT1 but not ANGPT2/4 is required for the formation of BM sinusoids. **A.** Tamoxifen intragastric (i.g.) administration and analysis scheme. **B.** Quantification of *Angpt1* transcript levels in lung tissues by qRT-PCR to examine gene deletion efficiency in *Angpt1^iKO;UBC+^* mice at P21 (*Control*: 1.00 ± 0.43, n=3; *Angpt1^iKO;UBC+^*: 0.22 ± 0.05, n=7, *P*=0.0857). **C.** Analysis of bone marrow sinusoidal morphogenesis and VEGFR3 /CAV1 expression in sinusoidal ECs of *Angpt1^iKO;UBC+^* and littermate control mice at P11. **D**. Quantification of VEGFR3 fluorescence relative to EMCN in diaphysis of *Angpt1^iKO;UBC+^* and littermate control mice at P11 (*Control*: 1.00 ± 0.12, n=5, *Angpt1^iKO;UBC+^*: 0.59 ± 0.23, n=5, P=0.0070). **E.** Western blot analysis of VEGFR3 in bone marrow and quantification (VEGFR3/beta-actin, *Control*: 1.00 ± 0.02, n=3; *Angpt1^iKO;UBC+^*: 0.50 ± 0.12, n=3, *P*=0.0018). **F**. Quantitative analysis of sinusoidal vessel diameter in the central diaphysis at P11 (*Control*: 20.14 ± 1.03 µm, n=5; *Angpt1^iKO;UBC+^*: 14.50 ± 0.84 µm, n=5, *P*<0.0001). **G-H**. Tamoxifen treatment and analysis scheme, and qRT-PCR analysis of *Angpt2* mRNA in lung tissues from *Angpt2^iKO;CDH5+^* and control mice at P7 (*Control*: 1.00 ± 0.39, n=9; *Angpt2^iKO;CDH5+^*: 0.37 ± 0.17, n=9, *P*<0.0001). **I**. Analysis of retinal vascularization by PECAM1 immunostaining and quantification of retinal vascularization index in *Angpt2^iKO;CDH5+^* mice and controls at P7 (ratio of vascularized area to total retinal area, *Control*: 1.00 ± 0.07, n=5; *Angpt2^iKO;CDH5+^*: 0.71 ± 0.05, n=6, *P*=0.0043). **J**. Analysis of bone marrow vascular network, VEGFR3 and CAV1 expression in BM sinusoidal vessels of *Angpt2^iKO;CDH5+^* and littermate control mice at P11. **K**. Analysis of ANGPT4 protein in lungs of *Angpt4*-deficient mice and littermate controls (2 month-old). **L** and **M.** Analysis of bone marrow vascular network, VEGFR3, EMCN, PECAM1 and αSMA expression in BM sinusoidal ECs of *Angpt4^-/-^* and littermate control mice at P21. Scale bar: 100 μm in C, J, L, M; 25 μm in I.

In contrast, the postnatal deletion of endothelial *Angpt2* did not produce an obvious effect on BM sinusoidal development (**Fig. 5G-J**). Shown in **Fig. 5G** is the scheme for *Angpt2* deletion by the intragastric administration of tamoxifen. The deletion efficiency of *Angpt2* was examined by the quantitative RT-PCR (**Fig. 5H**), and also confirmed by the suppression of retinal vascularization as shown in **Fig. 5I**. There was no obvious change with the BM sinusoidal network as well as the expression of sinusoidal VEGFR3 and CAV1 by the immunostaining (**Fig. 5J**). Furthermore, loss of ANGPT4 did not affect the bone marrow sinusoidal development and also the expression of VEGFR3 in SECs (**Fig. 5K-M**).

### Reciprocal regulation between COUP-TFII and TIE2 in BM sinusoidal development

Our previous study showed that the TIE2-mediated AKT activation was crucial for the specification of veins via regulating the protein stability of the venous fate determinant COUP-TFII [11]. On the other hand, COUP-TFII drives the expression of TIE2 ligand angiopoietin-1 (ANGPT1) in pericytes [14]. To investigate the requirement of COUP-TFII in murine BM sinusoidal morphogenesis, we generated a conditional knockout mouse model targeting the endothelial *Nr2f2* (*Nr2f2^Flox/Flox^*; *Cdh5-Cre^ERT2^*, named *Nr2f2^iKO;CDH5+^*). The scheme for the deletion of endothelial *Nr2f2* by the intragastric administration of tamoxifen is shown in **Fig. 6A**. COUP-TFII insufficiency resulted in reduced expression of VEGFR3 when analyzed 48 hours after the induced gene deletion (**Fig. 6B-C**). When analyzed at the stage of 96 hours, the disruption of the BM sinusoidal development in the *Nr2f2^iKO;CDH5+^*mutants was accompanied with a massive increase of angiogenesis, showing a decrease of VEGFR3 but an increase of CAV1 in BM blood vessels (**Fig. 6D**).

**Fig. 6.**
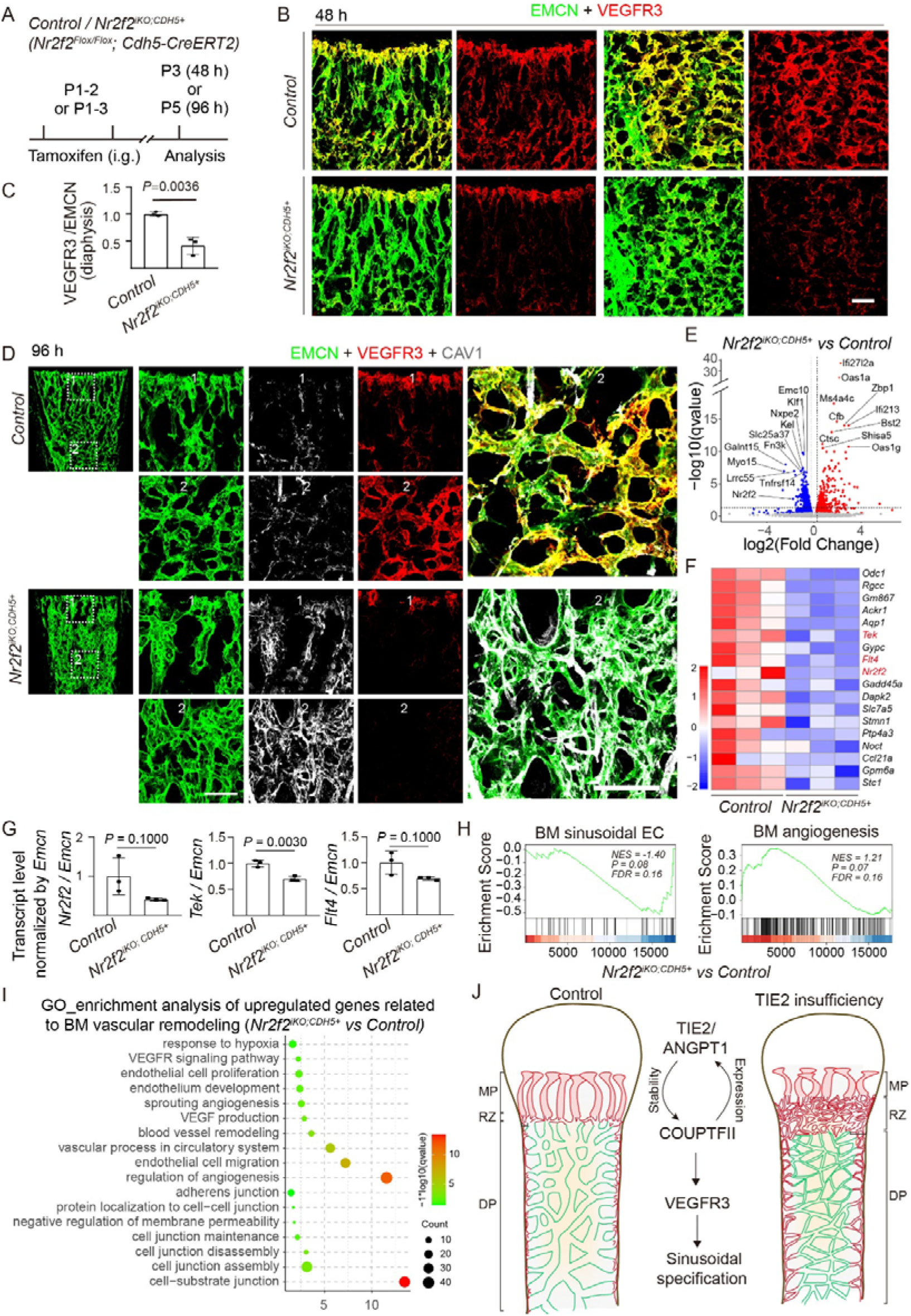
COUP-TFII regulates endothelial TIE2 and VEGFR3 in the BM sinusoidal development. **A.** Tamoxifen intragastric (i.g.) administration and analysis scheme. **B**-**D**. Analysis of bone marrow sinusoidal vessels, and VEGFR3 /CAV1 expression in sinusoidal ECs of *Nr2f2^iKO;CDH5+^* and littermate control mice at 48 h (B) and 96 h (D). Quantification of VEGFR3 fluorescence area relative to EMCN in the central diaphysis at 48 h (C; *Control*: 1.00 ± 0.04, n=3; *Nr2f2^iKO;CDH5+^*: 0.42 ± 0.16, n=3, *P*=0.0036). **E**-**I**. RNA-seq analysis of femur tissues from *Nr2f2^iKO;CDH5+^*and control mice at 96 h. Volcano plot showing differentially regulated genes (DEGs, P-value ≤ 0.05, |Log2Foldchange| > 0, E). Heatmap displaying subsets of downregulated sinusoidal endothelial genes (*P*-value ≤ 0.05, F). Quantitative expression analysis of sinusoidal endothelial genes (*Nr2f2*, *Tek*, *Flt4*) at 96 h (G). GSEA analysis showed that the genes involved in sinusoidal development were significantly downregulated after the endothelial deletion of *Nr2f2*, accompanied by the upregulation of angiogenesis-related genes (H). GO analysis further confirmed that genes related to hypoxia, angiogenesis and endothelial cell junction remodeling were significantly upregulated (I). **J**. Schematic illustration showing that ANGPT1/TIE2 and COUP-TFII form a reciprocal regulatory loop to coordinate BM angiogenic growth and sinusoidal specification via regulating the expression of VEGFR3. MP: Metaphysis, RZ: Remodeling zone, DP: Diaphysis. Scale bar: 100 μm.

To explore the mechanism underlying COUP-TFII in BM sinusoidal formation, we performed the RNA sequencing (RNA-seq) analysis of femur tissues after the induced deletion of endothelial *Nr2f2*. COUP-TFII insufficiency led to a significant decrease of gene expression related to the sinusoidal development. The altered expression of sinusoidal EC genes was shown by the volcano plot and heatmap (**Fig. 6E-F**). Shown in **Fig. 6G** is the transcript levels of key sinusoidal EC regulators, including *Tek* and *Flt4*, after the normalization by *Emcn*. Consistently, pathways related to sinusoidal development were downregulated, while the angiogenesis pathway was upregulated by the GSEA analysis using the custom gene sets as described above (**Fig. 6H**). This was further supported by the GO analysis showing that GO terms enriched for the downregulated genes include biological processes related to the sinusoidal morphogenesis, while pathways related to the hypoxia and angiogenic growth were upregulated in the bone marrow of *Nr2f2* mutants (**Fig. 6I**). COUP-TFII has also been shown to regulate the sinusoidal endothelial cell identity in zebrafish [37]. Taken together, these findings suggest that ANGPT1/TIE2 form a positive feedback loop with COUP-TFII to coordinate the angiogenic growth and sinusoidal specification via the regulation of VEGFR3 expression (**Fig. 6J**).

## Discussion

Sinusoidal morphogenesis in the bone marrow (BM) involves angiogenic expansion followed by remodeling into a discontinuous fenestrated network. VEGFR3 (*Flt4*) participates in the regulation of BM sinusoidal development [7], yet the upstream regulatory network remains inadequately characterized. We show in this study that the induced excision of endothelial TIE2 or its ligand ANGPT1 disrupted the process of vascular remodeling to form the mature bone marrow sinusoids, displaying a phenotype of sinusoidal capillarization. We further showed that TIE2 insufficiency led to a dramatic decrease of key sinusoidal regulators including VEGFR3. This is supported by the finding that the induced deletion of endothelial *Nr2f2* disrupted BM sinusoidal morphogenesis, displaying also decreased expression of TIE2 and VEGFR3. Findings from this study imply that the ANGPT1-TIE2 and COUPTFII form a positive feedback loop in the coordination of BM angiogenic expansion and sinusoidal specification via VEGFR3.

During embryonic development in mammals, the expression of VEGFR3 is detected in venous and capillary endothelial cells, but relatively low in arterial ECs [16]. At later stages of embryogenesis, VEGFR3 is mainly expressed by lymphatic ECs, which mediates a crucial pathway in lymphangiogenesis [16, 17]. Interestingly, VEGFR3 is also expressed by a proportion of blood vascular endothelial cells (BECs) including sinusoids. In addition to its critical role in lymphatic development, activation of endothelial VEGFR3 by VEGFC induces the VE-cadherin endocytosis via SRC-mediated phosphorylation, which is required for sinusoidal formation in both liver and bone marrow [7]. An interesting question is how endothelial TIE2 insufficiency affects VEGFR3 levels in BM sinusoidal vessels. TIE1/TIE2-mediated AKT activation has been shown to regulate the surface expression of lymphatic VEGFR3, therefore participating in the regulation of lymphangiogenesis [38]. We showed in this study that TIE2 protein expression was nearly undetectable by immunostaining 24 hours following induced endothelial *Tek* excision while there was no obvious change of the endothelial VEGFR3 in BM sinusoids. A significant decrease of VEGFR3 was detected by 48 hours upon the endothelial TIE2 insufficiency. Our previous study showed that TIE2-mediated activation of PI3K/AKT pathway is crucial for the protein stability of the transcription factor COUP-TFII [11]. COUP-TFII is a venous EC fate determining factor and participates in the regulation of atria and blood vascular development [13, 39]. COUP-TFII has also been shown to regulate the expression of NRP2 and VEGFR3 in lymphatic endothelial cells [15]. Therefore, the alteration of VEGFR3 expression by the TIE2 insufficiency may also occur at the transcription level, which result from the decrease of COUP-TFII. This was supported by the RNA-seq analysis, showing that the induced deletion of endothelial *Tek* resulted in a significant decrease of sinusoidal *Flt4* (encoding VEGFR3) within 48 hours. This was further supported by the finding that the endothelial-specific *Nr2f2* disrupted the sinusoidal vessel formation in bone marrow, showing a decrease of sinusoidal VEGFR3. Consistently, COUP-TFII was shown to regulate BM sinusoidal vessel formation in Zebrafish [37].

In addition, ANGPT1 and TIE2-mediated signals are vital for the specification of venous EC identity [11, 29, 40]. To exclude the possibility that the disruption of sinusoidal vessels resulted indirectly from the defective BM central veins, we performed studies with the induced *Tek* deletion at later stages with no obvious effect on veins. Furthermore, we have also generated a new Cre mouse line with its expression under the promoter of VEGFR3, which is mainly expressed in sinusoidal ECs in bone marrow. We provided evidences that the BM-SEC specific *Tek* deletion suppressed bone marrow sinusoidal vessel formation in the neonatal mice but with little effect on BM central veins. Consistently, the induced deletion of TIE2 ligand ANGPT1 produced a similar BM sinusoidal vascular phenotype. However, the endothelial ANGPT2 insufficiency or ANGPT4 deficiency did not produce an obvious effect on the bone marrow sinusoidal morphogenesis although they were shown to participate in the regulation of angiogenesis [41] and/or retinal vein formation [42].

Interestingly, the endothelial TIE2 insufficiency did not produce an obvious change of VEGFR3 expression in hepatic sinusoidal vessels at embryonic or postnatal stages. This suggests that the regulatory mechanism differs in bone marrow and liver sinusoidal vessel formation. Indeed, hepatic sinusoidal endothelial cells (SECs) are early specialized structures and are maintained by GATA4-mediated gene regulation. The GATA4 deficiency induces the liver sinusoidal capillarization, leading to hepatic hypoplasia [43, 44]. In contrast, GATA4 was not detected in bone marrow sinusoids as shown by the RNA-seq analysis. However, we found that GATA2 was upregulated after the induced excision of endothelial *Tek*. GATA2 has been shown to regulates lymphatic EC junctional integrity, lymphovenous valves [45] and the endothelial-to-hematopoietic transition [46]. Further investigation is required to elucidate how GATA2 and other downstream components of the TIE2 pathway regulate bone marrow sinusoidal specification.

In conclusion, the endothelial TIE2 and its ligand ANGPT1 are crucial for the bone marrow sinusoidal morphogenesis. This may rely on the TIE2-mediated activation of PI3K-AKT signaling, which regulates the COUP-TFII protein stability [11]. On the other hand, COUP-TFII drives the transcription of the TIE2 ligand angiopoietin-1 (ANGPT1) in pericytes [14], as well as the expression of VEGFR3 as shown in this study and also by other researchers [15]. These findings suggest a positive feedback loop between TIE2 and COUP-TFII in the regulation of BM sinusoidal morphogenesis via regulating VEGFR3 expression in SECs.

## Acknowledgement

We thank staff in the Animal facility of Soochow University for technical assistance. This work was supported by grants from the National Natural Science Foundation of China (82470518, 82401544), China Postdoctoral Science Foundation (CPSF, 2024M762300), the National Key R&D Program of China (2021YFA0805000), Cyrus Tang Foundation (CTJC25001), the Natural Science Foundation of Jiangsu Province (BK20240786), the Project of State Key Laboratory of Radiation Medicine and Protection (No. GZN120 20 02), and the Priority Academic Program Development of Jiangsu Higher Education Institutions.

## Reference

1. Augustin, H.G. and G.Y. Koh, Organotypic vasculature: From descriptive heterogeneity to functional pathophysiology. Science, 2017. 357(6353).

2. Lucas, D., Structural organization of the bone marrow and its role in hematopoiesis. Curr Opin Hematol, 2021. 28(1): p. 36–42.

3. Langen, U.H., et al., Cell-matrix signals specify bone endothelial cells during developmental osteogenesis. Nat Cell Biol, 2017. 19(3): p. 189–201.

4. Liu, Q., et al., Genetic targeting of sprouting angiogenesis using Apln-CreER. Nat Commun, 2015. 6: p. 6020.

5. Kusumbe, A.P., S.K. Ramasamy, and R.H. Adams, Coupling of angiogenesis and osteogenesis by a specific vessel subtype in bone. Nature, 2014. 507(7492): p. 323–8.

6. Weinhaus, B., et al., Differential regulation of fetal bone marrow and liver hematopoiesis by yolk-sac-derived myeloid cells. Nat Commun, 2025. 16(1): p. 4427.

7. Sung, D.C., et al., Sinusoidal and lymphatic vessel growth is controlled by reciprocal VEGF-C-CDH5 inhibition. Nat Cardiovasc Res, 2022. 1(11): p. 1006–1021.

8. Ramasamy, S.K., et al., Endothelial Notch activity promotes angiogenesis and osteogenesis in bone. Nature, 2014. 507(7492): p. 376–80.

9. Schipani, E., et al., Regulation of Bone Marrow Angiogenesis by Osteoblasts during Bone Development and Homeostasis. Front Endocrinol (Lausanne), 2013. 4: p. 85.

10. Augustin, H.G., et al., Control of vascular morphogenesis and homeostasis through the angiopoietin-Tie system. Nat Rev Mol Cell Biol, 2009. 10(3): p. 165–77.

11. Chu, M., et al., Angiopoietin receptor Tie2 is required for vein specification and maintenance via regulating COUP-TFII. Elife, 2016. 5:e21032.

12. Chen, X., et al., COUP-TFII is a major regulator of cell cycle and Notch signaling pathways. Mol Endocrinol, 2012. 26(8): p. 1268–77.

13. You, L.R., et al., Suppression of Notch signalling by the COUP-TFII transcription factor regulates vein identity. Nature, 2005. 435(7038): p. 98–104.

14. Qin, J., et al., COUP-TFII regulates tumor growth and metastasis by modulating tumor angiogenesis. Proc Natl Acad Sci U S A, 2010. 107(8): p. 3687–92.

15. Lin, F.J., et al., Direct transcriptional regulation of neuropilin-2 by COUP-TFII modulates multiple steps in murine lymphatic vessel development. J Clin Invest, 2010. 120(5): p. 1694–707.

16. Zhang, L., et al., VEGFR-3 ligand-binding and kinase activity are required for lymphangiogenesis but not for angiogenesis. Cell Res, 2010. 20(12): p. 1319–31.

17. Makinen, T., et al., Inhibition of lymphangiogenesis with resulting lymphedema in transgenic mice expressing soluble VEGF receptor-3. Nat Med, 2001. 7(2): p. 199–205.

18. Sivaraj, K.K., et al., YAP1 and TAZ negatively control bone angiogenesis by limiting hypoxia-inducible factor signaling in endothelial cells. Elife, 2020. 9.

19. Shao, L., et al., A Tie2-Notch1 signaling axis regulates regeneration of the endothelial bone marrow niche. Haematologica, 2019. 104(11): p. 2164–2177.

20. Kopp, H.G., et al., Tie2 activation contributes to hemangiogenic regeneration after myelosuppression. Blood, 2005. 106(2): p. 505–13.

21. Shen, B., et al., Genetic Dissection of Tie Pathway in Mouse Lymphatic Maturation and Valve Development. Arteriosclerosis, thrombosis, and vascular biology, 2014. 34.

22. Farley, F.W., et al., Widespread recombinase expression using FLPeR (flipper) mice. Genesis, 2000. 28(3-4): p. 106–10.

23. Ruzankina, Y., et al., Deletion of the developmentally essential gene ATR in adult mice leads to age-related phenotypes and stem cell loss. Cell Stem Cell, 2007. 1(1): p. 113–26.

24. Okabe, K., et al., Neurons limit angiogenesis by titrating VEGF in retina. Cell, 2014. 159(3): p. 584–96.

25. Nagy, A., et al., Derivation of completely cell culture-derived mice from early-passage embryonic stem cells. Proc Natl Acad Sci U S A, 1993. 90(18): p. 8424–8.

26. Takamoto, N., et al., COUP-TFII is essential for radial and anteroposterior patterning of the stomach. Development, 2005. 132(9): p. 2179–89.

27. Cao, X., et al., A Genetically Engineered Mouse Model of Venous Anomaly and Retinal Angioma-like Vascular Malformation. Bio Protoc, 2021. 11(15): p. e4117.

28. Pitulescu, M.E., et al., Inducible gene targeting in the neonatal vasculature and analysis of retinal angiogenesis in mice. Nat Protoc, 2010. 5(9): p. 1518–34.

29. Cao, X., et al., Endothelial TIE1 Restricts Angiogenic Sprouting to Coordinate Vein Assembly in Synergy With Its Homologue TIE2. Arterioscler Thromb Vasc Biol, 2023. 43(8): p. e323–e338.

30. Bandyopadhyay, S., et al., Mapping the cellular biogeography of human bone marrow niches using single-cell transcriptomics and proteomic imaging. Cell, 2024. 187(12): p. 3120–3140 e29.

31. Kalucka, J., et al., Single-Cell Transcriptome Atlas of Murine Endothelial Cells. Cell, 2020. 180(4): p. 764–779 e20.

32. Liberzon, A., et al., The Molecular Signatures Database (MSigDB) hallmark gene set collection. Cell Syst, 2015. 1(6): p. 417–425.

33. Hooper, A.T., et al., Engraftment and reconstitution of hematopoiesis is dependent on VEGFR2-mediated regeneration of sinusoidal endothelial cells. Cell Stem Cell, 2009. 4(3): p. 263–74.

34. Mohanakrishnan, V., et al., Specialized post-arterial capillaries facilitate adult bone remodelling. Nat Cell Biol, 2024. 26(12): p. 2020–2034.

35. Liu, Y., et al., A specialized bone marrow microenvironment for fetal haematopoiesis. Nat Commun, 2022. 13(1): p. 1327.

36. Poulos, M.G., et al., Complementary and Inducible creER(T2) Mouse Models for Functional Evaluation of Endothelial Cell Subtypes in the Bone Marrow. Stem Cell Rev Rep, 2024. 20(4): p. 1135–1149.

37. Hagedorn, E.J., et al., Transcription factor induction of vascular blood stem cell niches in vivo. Dev Cell, 2023. 58(12): p. 1037–1051 e4.

38. Korhonen, E.A., et al., Lymphangiogenesis requires Ang2/Tie/PI3K signaling for VEGFR3 cell-surface expression. J Clin Invest, 2022. 132(15).

39. Wu, S.P., et al., Atrial identity is determined by a COUP-TFII regulatory network. Dev Cell, 2013. 25(4): p. 417–26.

40. Arita, Y., et al., Myocardium-derived angiopoietin-1 is essential for coronary vein formation in the developing heart. Nat Commun, 2014. 5: p. 4552.

41. Maisonpierre, P.C., et al., Angiopoietin-2, a natural antagonist for Tie2 that disrupts in vivo angiogenesis. Science, 1997. 277(5322): p. 55–60.

42. Elamaa, H., et al., Angiopoietin-4-dependent venous maturation and fluid drainage in the peripheral retina. Elife, 2018. 7.

43. Géraud, C., et al., GATA4-dependent organ-specific endothelial differentiation controls liver development and embryonic hematopoiesis. The Journal of Clinical Investigation, 2017. 127(3): p. 1099–1114.

44. Griffin, C.T. and S. Gao, Building discontinuous liver sinusoidal vessels. J Clin Invest, 2017. 127(3): p. 790–792.

45. Mahamud, M.R., et al., GATA2 controls lymphatic endothelial cell junctional integrity and lymphovenous valve morphogenesis through miR-126. Development, 2019. 146(21).

46. Koyunlar, C., et al., Gata2-regulated Gfi1b expression controls endothelial programming during endothelial-to-hematopoietic transition. Blood Adv, 2023. 7(10): p. 2082–2093.

