## Supplemental Fig. 1-5 for "Specification of bone marrow sinusoids requires TIE2-mediated positive feedback involving COUPTFII and VEGFR3"

Short title: TIE2 in bone marrow sinusoidal specification

Key words: sinusoidal morphogenesis, bone marrow, angiopoietin recetpor TIE2, COUPTFII, VEGFR3, genetically modified mouse model

[*]Correspondence should be addressed to Dr. Yulong He

### Supplemental figures and figure legends

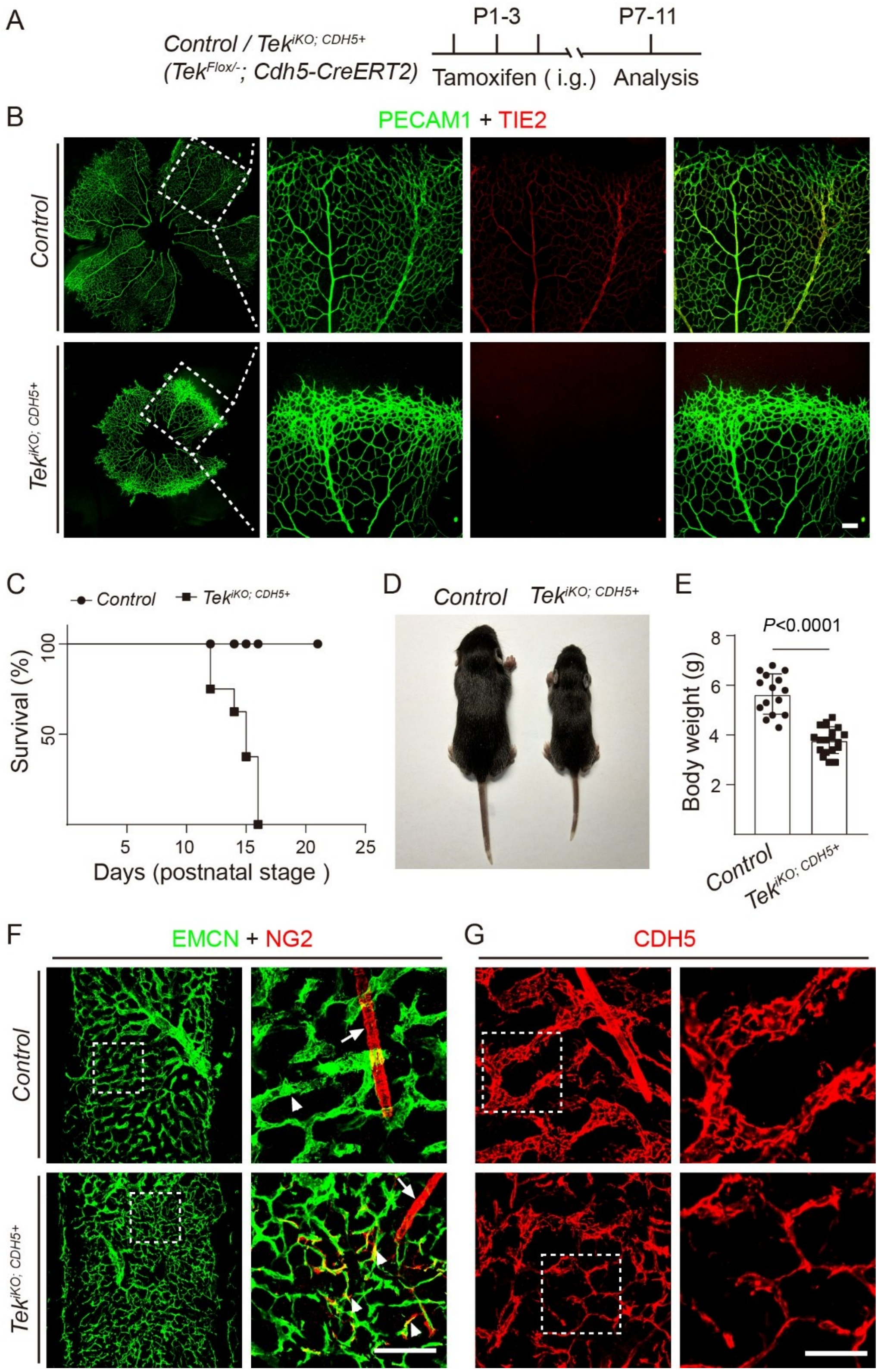


**Supplemental Fig. 1 TIE2 insufficiency leads to the abnormal angiogenesis and mural cell recruitment with BM sinusoidal vessels. A.** Tamoxifen intragastric (i.g.) administration and analysis scheme. **B.** Analysis of endothelial *Tek* deletion by immunostaining for PECAM1 and TIE2 in retinal vessels at P7. **C.** Survival curve of *Tek*$^{iKO;CDH5+}$ and littermate control mice after the endothelial *Tek* deletion (n=8 per group, *P*=0.0006, Log-rank (Mantel-Cox) test). **D.** Representative image of *Tek*$^{iKO;CDH5+}$ and littermate control mice at P11. **E.** Quantification of body weight of *Tek*$^{iKO;CDH5+}$ mutants and the littermate controls (*Control*: 5.65 ± 0.81 g, n=15; *Tek*$^{iKO;CDH5+}$: 3.79 ± 0.54 g, n=19, *P*<0.0001). **F** and **G.** Analysis of abnormal mural cell coverage with BM sinusoidal vessels (NG2$^{+}$, F) and endothelial adherens junctions (CDH5, G) in *Tek*$^{iKO;CDH5+}$ and littermate control mice at P11. Arrows indicate artery, arrowheads indicate sinusoidal vessels. Scale bar: 40 μm in G, other scale bar: 100 μm.

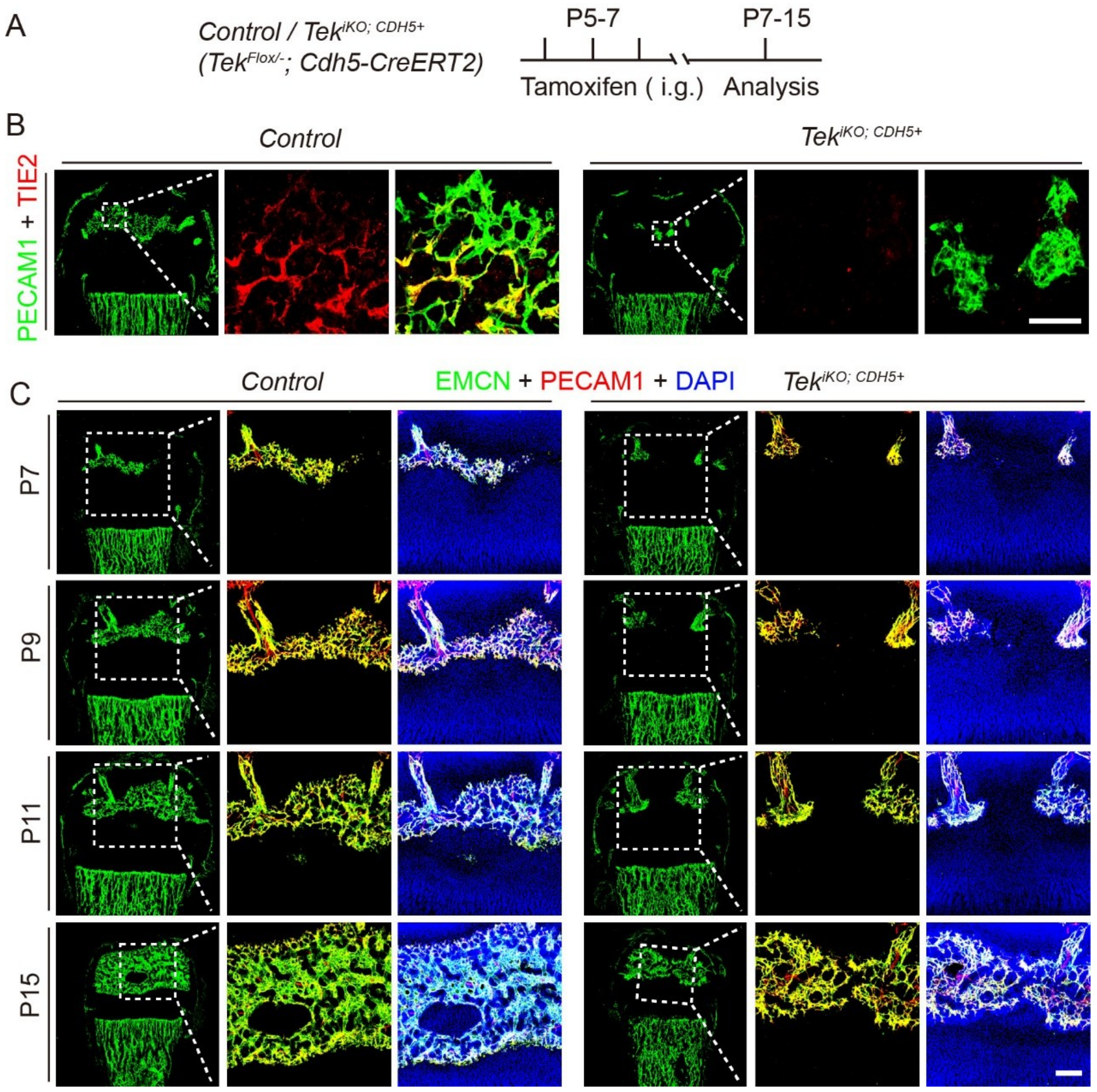


**Supplemental Fig. 2 TIE2 insufficiency delays the vascularization of secondary ossification center in the femur. A.** Tamoxifen intragastric (i.g.) administration and analysis scheme. **B-C.** Analysis of endothelial *Tek* deletion and vascular formation in the BM secondary ossification center of *Tek*$^{iKO;CDH5+}$ and littermate control mice by immunostaining for TIE2, PECAM1 and EMCN at different time points following tamoxifen treatment initiated at P5-P7. Scale bar:100 μm.

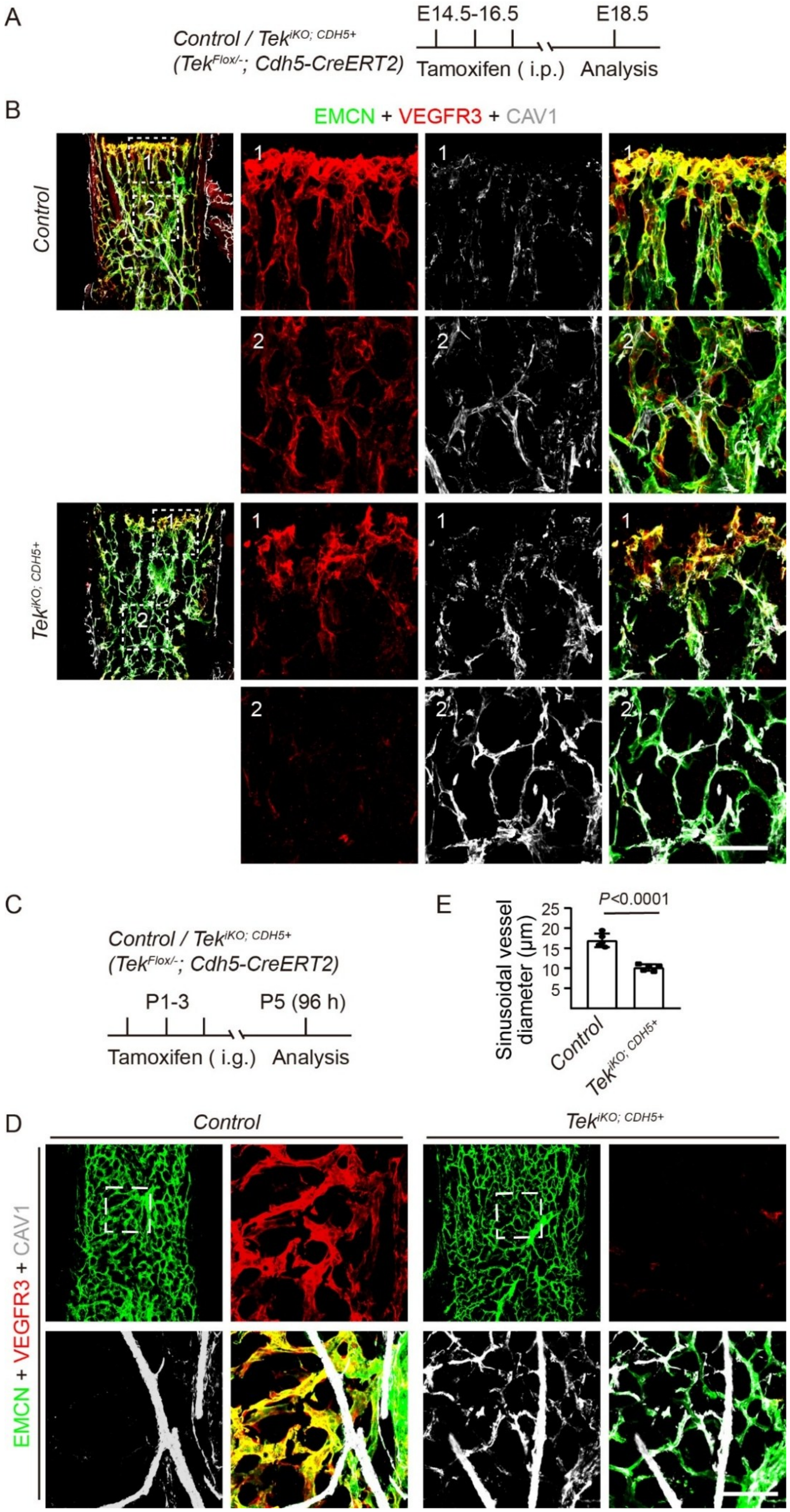
A
Control / Tek$^{iKO; CDH5+}$
(Tek$^{Flox/-}$; Cdh5-CreERT2)
E14.5-16.5
E18.5
Tamoxifen ( i.p.)
Analysis
B
EMCN + VEGFR3 + CAV1
Control
Tek$^{iKO; CDH5+}$
C
Control / Tek$^{iKO; CDH5+}$
(Tek$^{Flox/-}$; Cdh5-CreERT2)
P1-3
P5 (96 h)
Tamoxifen ( i.g.)
Analysis
E
P<0.0001
Sinusoidal vessel diameter (μm)
Control
Tek$^{iKO; CDH5+}$
D
Control
Tek$^{iKO; CDH5+}$
EMCN + VEGFR3 + CAV1

**Supplemental Fig. 3 Disruption of BM sinusoidal morphogenesis 96 hours after the induced *Tek* deletion at embryonic and postnatal stages. A** and **C**. Tamoxifen intraperitoneal (i.p., A) and intragastric (i.g., C) administration and analysis scheme. **B** and **D**. Analysis of bone marrow sinusoidal vessels, VEGFR3 and CAV1 expression in sinusoidal ECs of $Tek^{iKO;CDH5+}$ and littermate control mice at E18.5 (B) and P5 (D). **E**. Quantitative analysis of sinusoidal vessel diameter in the diaphysis at P5 (*Control*: 16.96 ± 1.70 μm, n=5; $Tek^{iKO;CDH5+}$: 10.19 ± 0.80 μm, n=5, $P<0.0001$). CV: central vein. Scale bar:100 μm.

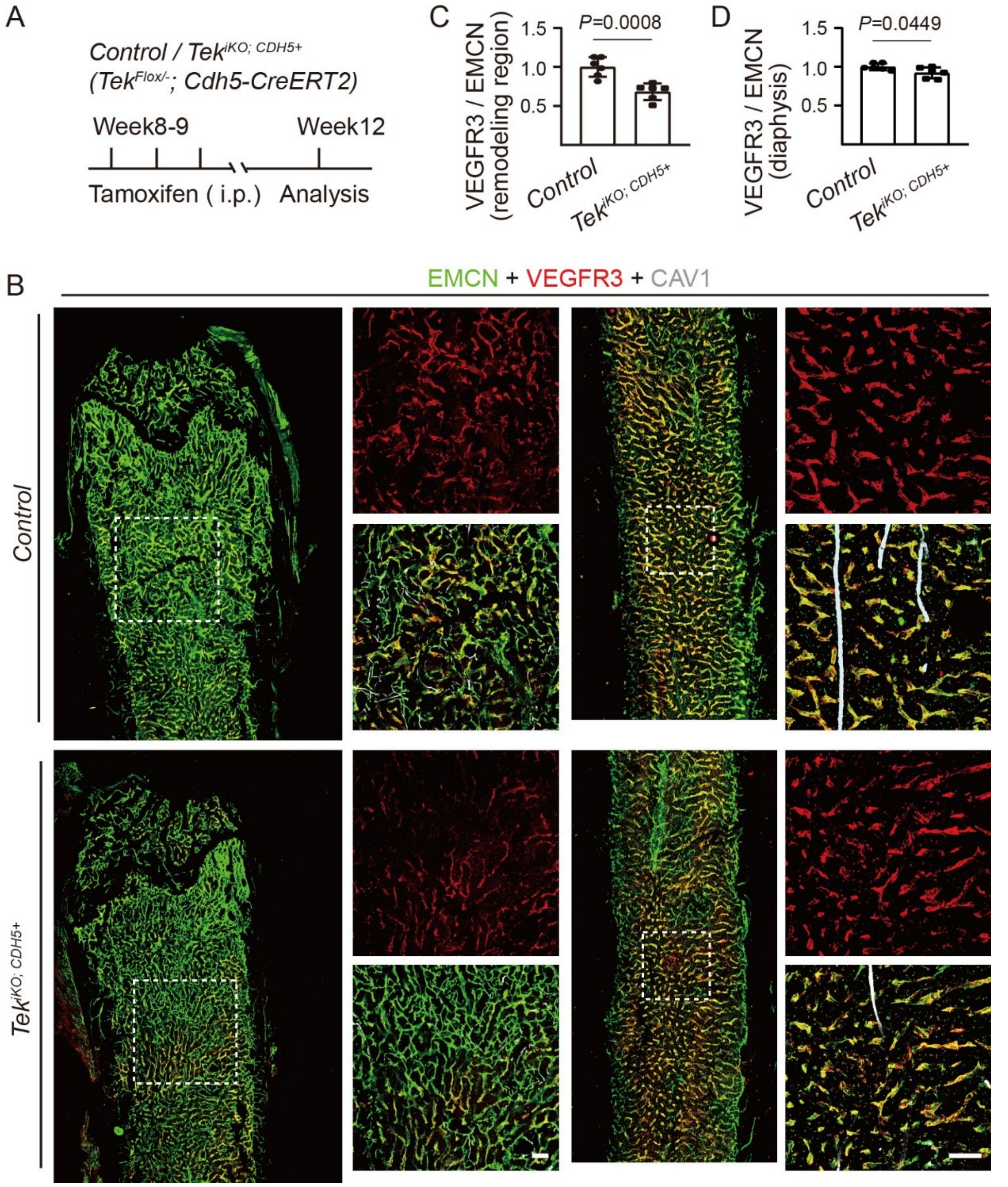


**Supplemental Fig. 4 Requirement of TIE2 in the maintenance of bone marrow sinusoids at the adult stage. A**. Tamoxifen intraperitoneal (i.p.) administration and analysis scheme. **B**. Analysis of bone marrow sinusoidal vessels, VEGFR3 and CAV1 expression in sinusoidal ECs of *Tek*$^{iKO;CDH5+}$ and littermate control mice after the induced gene deletion at adult stages (Week12). **C** and **D**. Quantification of VEGFR3 fluorescence intensity relative to EMCN in BM transition zone (*Control*: 1.00 ± 0.13, n=6; *Tek*$^{iKO;CDH5+}$: 0.68 ± 0.11, n=6, *P*=0.0008), and maturation zone of diaphysis at postnatal week 12 (*Control*: 1.00 ± 0.04, n=6; *Tek*$^{iKO;CDH5+}$: 0.92 ± 0.07, n=6, *P*=0.0449). Scale bar:100 μm.

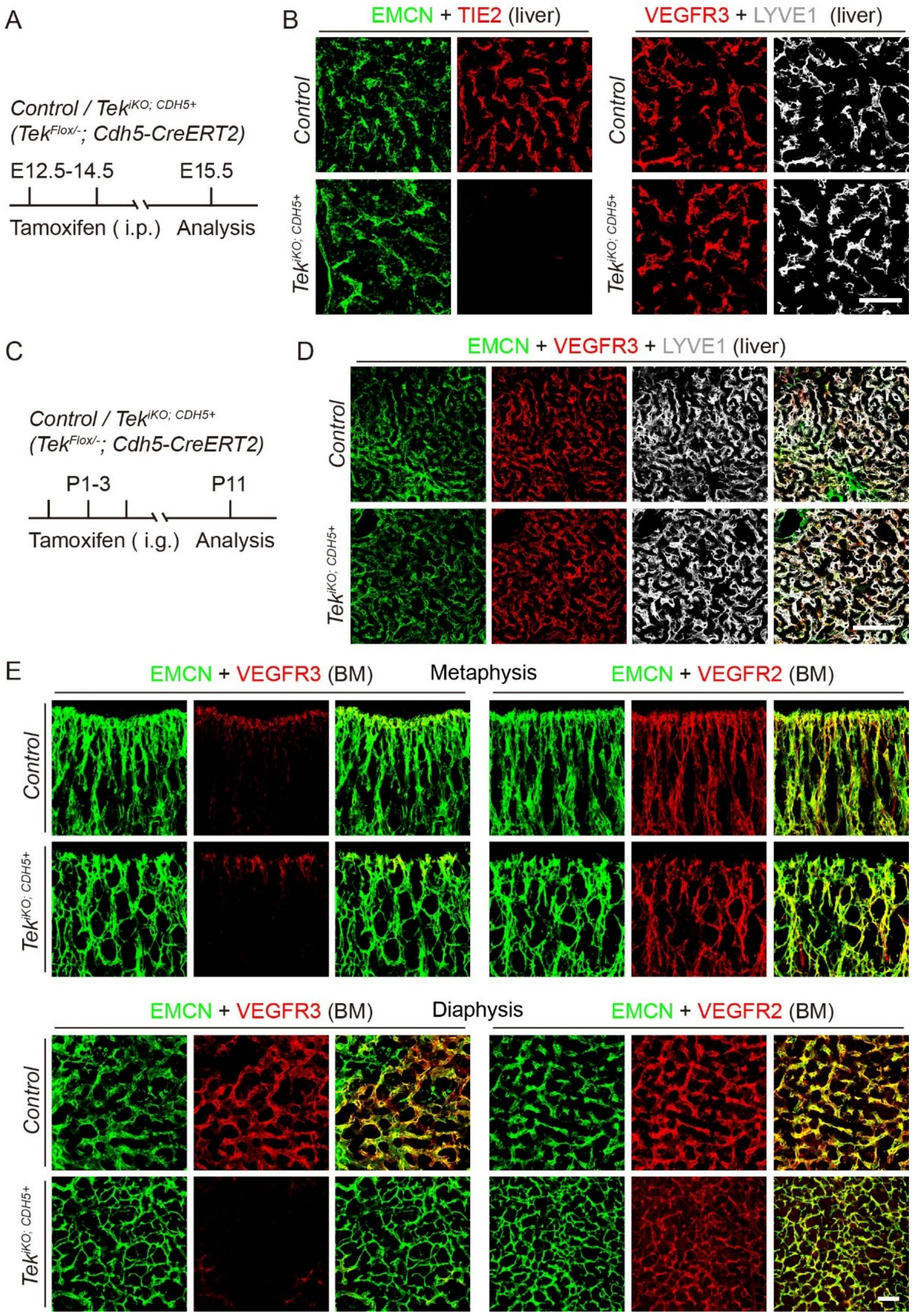


**Supplemental Fig. 5 Comparative analysis of sinusoidal VEGFR2 and VEGFR3 in the liver and bone marrow after the induced *Tek* deletion. A-D**. Tamoxifen treatment and analysis schemes at embryonic stages (A) and postnatal stages (C). Immunostaining analysis for TIE2 and VEGFR3 in the hepatic sinusoidal endothelial cells (positive for EMCN and LYVE1) in the *Tek*$^{iKO;CDH5+}$ mutants and littermate controls at E15.5 (B) and P11 (D). **E**. Comparative analysis of sinusoidal VEGFR3 and

VEGFR2 in the metaphysis and diaphysis of femur tissues from the *Tek*$^{iKO;CDH5+}$ mutants and littermate controls at P11. Scale bar:100 μm.
